# Ecology drives the observed spectrum of hydrophobin protein diversity across Kingdom Fungi

**DOI:** 10.1101/2022.08.19.504535

**Authors:** Brian Lovett, Matt T. Kasson, Julie-Anne Gandier

**Affiliations:** Emerging Pests and Pathogens Research Unit USDA ARS, Ithaca, New York, United States of America; Division of Plant and Soil Sciences, West Virginia University, Morgantown, United States of America; The Centre of Excellence in Life-Inspired Hybrid Materials (LIBER), Department of Bioproducts and Biosystems, School of Chemical Engineering, Aalto University, Espoo, Finland

## Abstract

Hydrophobins mediate the interactions between fungi and the elements of their ecosystem via assembly at interfaces serving a wide range of diverse functions. As such, these proteins can be seen as a means by which fungi not only adapt to a pre-existing environment, but also actively participate in the construction of their own ecological niches. Through this lens, we provide an expansive hydrophobin survey across the ecological breadth of Kingdom Fungi and advance the view that hydrophobins are best defined as a generic molecular structure with shared core structural features that accommodate a remarkable diversity of amino acid sequences. We examine the relationship between hydrophobin sequences, fungus phylogeny, and associated ecology from 45 fungal proteomes predicted from genomes spanning eight phyla and more than 25 orders. To capture the full spectrum of the hydrophobin amino acid sequence space mapped by our study, we describe the family as a continuum of overlapping hidden Markov models (HMMs), each HMM representing clusters of sequence similarity spanning existing hydrophobin classes. Overall, our approach uncovered ecology as a major driver of hydrophobin diversification, further expanded the known hydrophobins beyond Dikarya, and uncovered evidence extending the possibilities for their function from exclusively extracellular to include intracellular. In addition, we identified novel core groups of cysteine-rich proteins whose conservation across fungi suggest they play key ecological roles. Together, our work offers an ontological framework that captures the diversity of hydrophobin amino acid sequences and highlights the need to revisit challenging fundamental questions regarding hydrophobins to achieve a mechanistic understanding of their function as emerging from assembly within an ecosystem.

## 2. Introduction

Hydrophobins are a family of non-catalytic small secreted cysteine-rich fungal proteins (SSCPs) best known for mediating interactions between fungi and their ecosystem via self-assembly into films that serve as interfaces between the fungus and the different elements of its ecosystem. In other words, fungi secrete hydrophobins to tailor the physicochemical properties of the fungal environment to support their particular lifestyle. First coined by (Rosenberg and Kjelleberg, 1986), the term hydrophobin evokes a single intuitive function: the formation of water-repellent “hydrophobic” coatings as first observed at the surface of the fungal spore (Beever and Dempsey, 1978). Since their discovery, however, a rich diversity of biological functions have been ascribed to members of this family well beyond conferring hydrophobicity (detailed in Table 1). In fact, one study of *Trichoderma* species revealed that hydrophobin coatings modulate the wettability of the spore surface to suit a particular dispersion strategy. That is to say the hydrophobin coating at the surface of water-dispersed spores increases wettability, while the coatings at the surface of aerially-dispersed spores decrease wettability (Cai et al., 2020). Furthermore, in *Aspergillus fumigatus*, the hydrophobin layer has also been shown to mask the spore from both innate and adaptive human immune systems upon infection (Aimanianda et al., 2009).

**Table 1.**
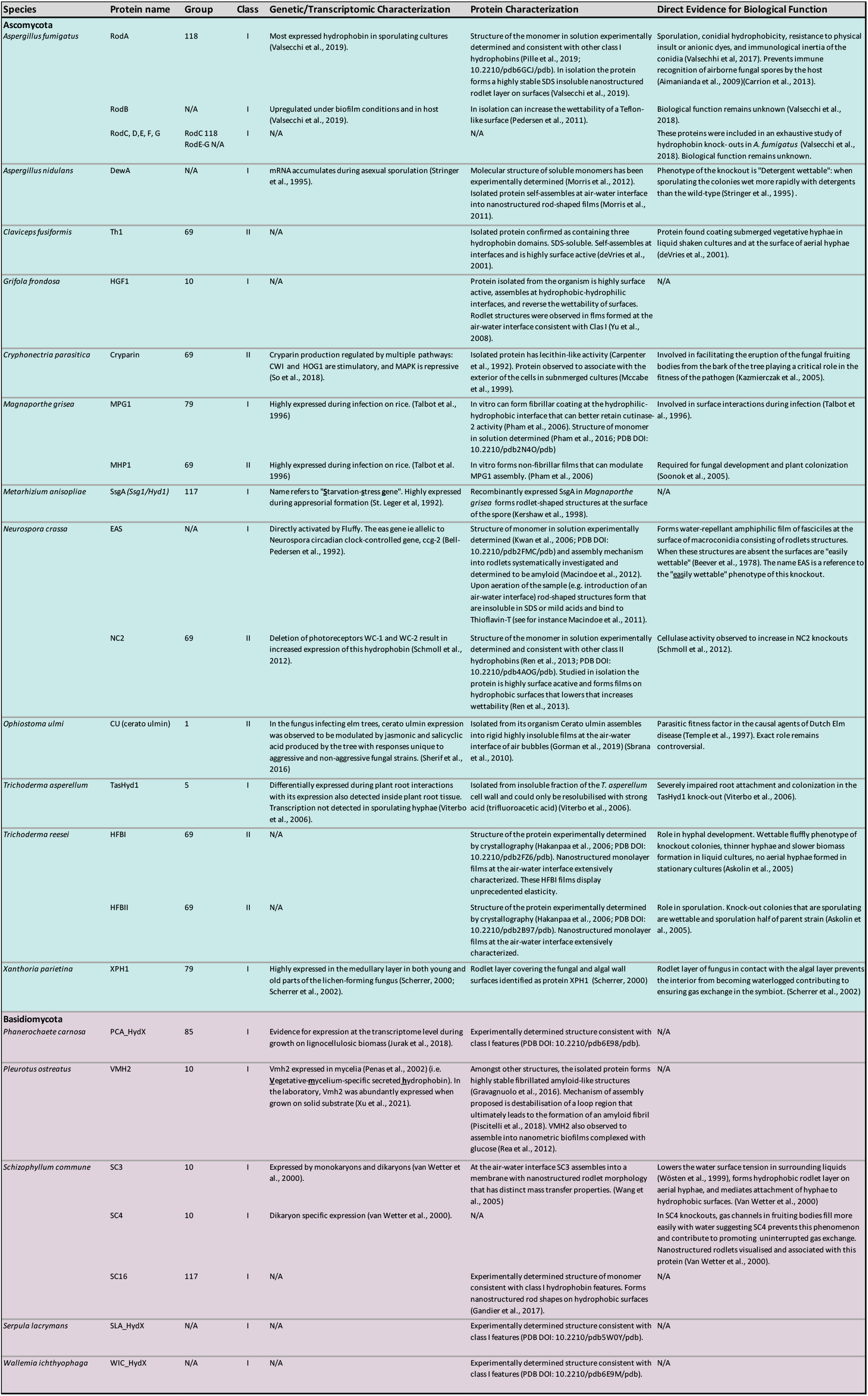
Functions associated with experimentally confirmed hydrophobin proteins (indicator sequences). Proteins whose biophysical properties were directly studied either in isolation or within their biological context. These sequences are referred to as “indicator sequences” throughout as they contribute to our definition of the hydrophobin amino acid sequence in addition to providing reference points for experimentally verified biophysical properties. Group number refers to the ontology developed in this study (**Section 5.9**).

Beyond forming a protective coat around spores, hydrophobins serve multiple functions critical to the biology and ecology of fungi in their environment. These functions are as varied as detecting plants by self-assembling into enzyme-recruiting coatings on the surface of leaves (Pham et al., 2016), preventing water logging of lichens by self-assembling at the interface of fungal and algal components (Scherrer et al., 2000), assembling to form fungal adhesion points on environmental surfaces (Wösten et al., 1994), and assembling at the air-liquid interface of submerged cultures reducing the surface tension to facilitate the emergence of aerial hyphae (Wessels et al., 1991a). Furthermore, fungal genomes typically encode suites of hydrophobins whose expression appear to be finely tuned, as has been shown in mycorrhizal fungi (Rineau et al., 2017), yet also often include functionally redundant hydrophobins (Shin et al., 2022). Given this complexity, the full range of biological functions to which hydrophobin proteins contribute has yet to be elucidated.

Despite the diversity of their biological function, members of the hydrophobin family are unified by a common theme: their function is a result of the protein’s assembly within a particular microenvironment, which the protein simultaneously responds to and modulates. As such hydrophobins can be seen as a means by which fungi not only adapt to a pre-existing environment, but are also active participants in the construction of their ecological niche. Hydrophobin function therefore cannot be considered exclusively as acting in isolation, but rather as emerging from the fluctuating properties of the ecosystem. This point of view—that hydrophobin function is shaped by biotic and abiotic context—is consistent with the pleiotropic effects observed in biological studies of hydrophobin knock-outs e.g (Terhem and van Kan, 2014).

Through this lens, we have conducted the most expansive hydrophobin survey to date across the ecological breadth of Kingdom Fungi. We build on the view that the hydrophobin family is best defined by a shared generic core structure that captures the remarkable sequence diversity accommodating their wide breadth of molecular and biological functions. As will be described in **Section 3** (defining the hydrophobin family) low sequence similarity (i.e., two members sharing as little as 10% identity in alignments) is a feature of the functional diversity of the family. As such, describing the sequence diversity of the full spectrum of these proteins poses a significant challenge as protein ontologies typically rely on similarity of amino acid sequences (or folded structures) to form discrete categories. Often, these discrete categories are used to infer protein function in a manner that is intertwined with questions of the family’s evolutionary origin through, for instance, the construction of phylogenetic trees. This exercise again relies on the conservation of the protein sequence over evolutionary time such that shared origins can be traced. Hydrophobin sequences considered together as a family however do not retain very much “evolutionary” information (Littlejohn et al., 2012; Mgbeahuruike et al., 2013; Plett et al., 2012). We hypothesize that the reason for this is that ecology is a major driver of diversification of the sequences.

In this paper we set aside evolutionary questions, to first catalog hydrophobin diversity across Kingdom Fungi, while highlighting how this diverse fungal hydrophobin repertoire relates to host phylogeny, lifestyle, and ecosystem. This step is necessary to develop fruitful questions that will further build a mechanistic understanding of hydrophobin assembly and function within their full context. That is, hydrophobin function emerges from assembly within an ecosystem. Hydrophobins don’t act in isolation: their sequences must be contextualised within their broader ecology to understand how they contribute to the construction of their ecological niches.

## 3. Defining the hydrophobin family: Multifunctional proteins that accommodate structural diversity

The term hydrophobin was first adopted by Wessels et al. (Wessels et al., 1991a, 1991b) to describe the protein products of previously characterized homologous genes (SC-1, SC-3, and SC-4) whose transcripts were abundantly and differentially detected during the development of *Schizophyllum commune* (Mulder and Wessels, 1986) (**Figure 1A**). Although challenging to isolate, the proteins were eventually found in the form of highly-insoluble assemblies associated with aerial hyphae and sexual fruiting bodies (Wessels et al., 1991a, 1991b). Given these proteins were secreted, associated with the cell wall as water-insoluble assemblies, and contained a significant proportion of hydrophobic amino acids, the proteins were collectively referred to as hydrophobins (Wessels et al., 1991a). In adopting the term hydrophobin, Wessels et al. established the hypothesis that these proteins assembled on fungal surfaces to increase their hydrophobicity. It would later be experimentally demonstrated that, among other functions, SC-3 indeed coats aerial hyphae with a hydrophobic “raincoat” (Wösten et al., 1994). This hydrophobic layer was also shown to mediate attachment to hydrophobic surfaces. SC-4 was found to coat the surface of gas channels within fruiting bodies, preventing water logging and contributing to uninterrupted gas exchange (Lugones et al., 1999). While distinct roles can be found for individual hydrophobin proteins within a particular organism, they typically play multiple roles and are functionally redundant to varying extents. For instance, both SC-3 and SC-4 can assemble at the air-water interface to reduce surface tension and facilitate the emergence of aerial structures from the liquid medium. The properties of these films, however, are not identical. The amphiphilic nature of the SC-3 film, for instance, is more pronounced than that of SC-4 (van Wetter et al., 2000).

**Figure 1.**
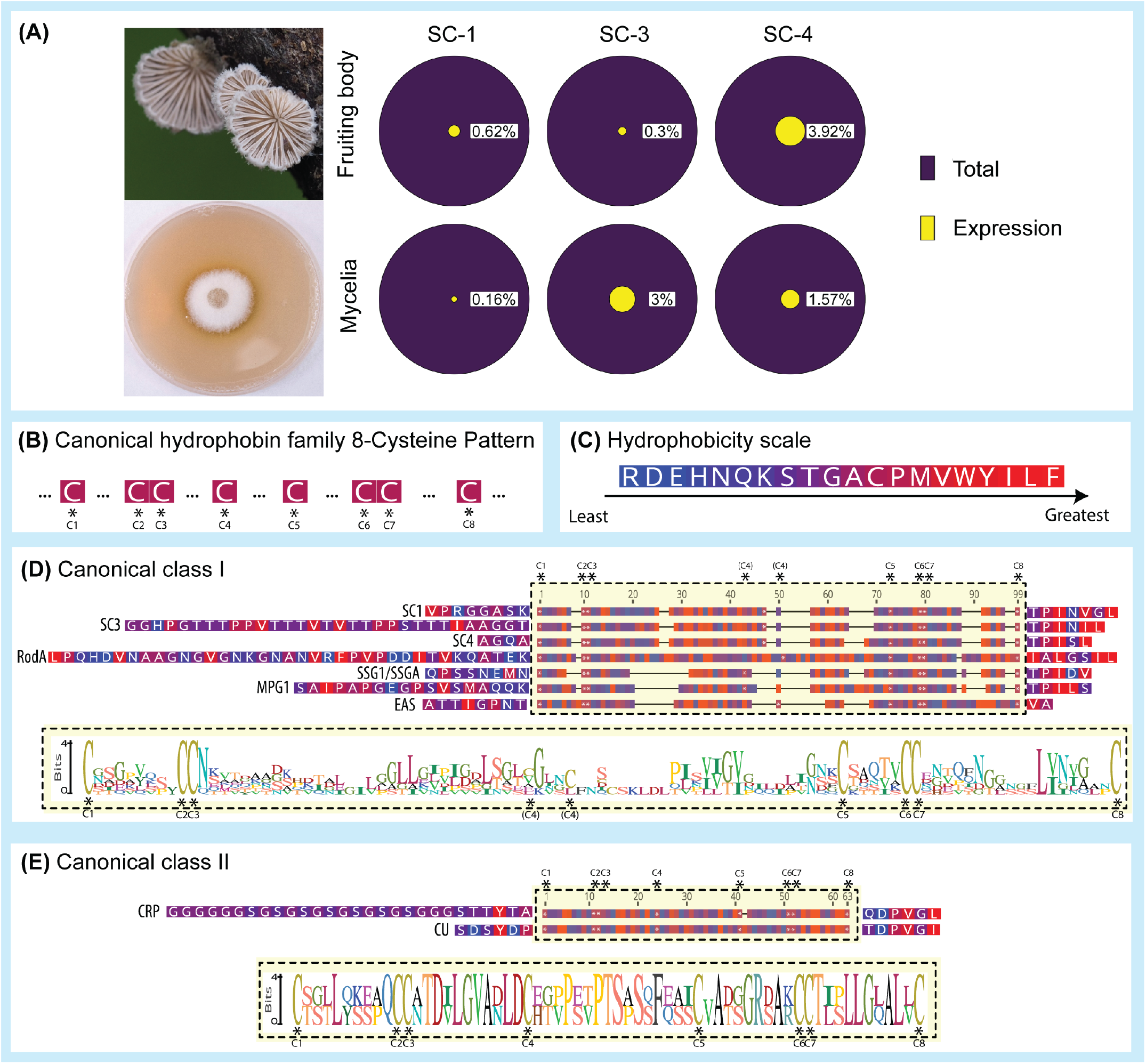
Founding members of the hydrophobin family shaped their canonical definition and subdivisions. (A) The first members of the hydrophobin family were the products of a group of homologous *Schizophyllum commune* genes with exceptional levels of transcription. The total proportion of mRNA that encodes hydrophobin proteins is shown for the fruiting body and mycelia summarized from (Mulder and Wessels, 1986). (B) The defining feature of these *S. commune* sequences are a strictly conserved pattern of eight cysteine residues (8-Cys pattern). This pattern was identified in sequences of predicted and confirmed proteins shown in (D) and (E). The amino acids of these sequences are coloured by (C) the physiological hydrophobicity of their amino acids (Black and Mould, 1991). The yellow boxes denote the region containing the characteristic 8-Cys pattern. Typically, it is this region (C1-C8) that is aligned when comparing hydrophobin sequences and is characterized by blocks of alternating hydrophobic and hydrophilic residues. These patterns, also referred to as hydropathy plots, are particularly well conserved in the regions surrounding the cysteines. The C1-C8 region forms the core of the folded protein (Figure 2). The hydrophobin family was further subdivided into two classes (I and II) based on the similarity of their hydropathy plots. Logos generated from the foundational sequences are displayed with their respective classes. Class II sequences (E) display conserved spacing in the 8-Cys pattern and significant amino acid similarity. Class I sequences (D), however, share limited sequence similarity besides the 8-Cys pattern when considered together. Class I hydropathy patterns are similar when considering the cysteine residues as points of reference. The sequences displayed are: (C) *Schizophyllum commune* SC1, SC3, SC4; *Aspergillus fumigatus* RodA; *Metarhizium anisopliae* SSG1/SSGA; *Magnaporthe grisea* MPG1; *Neurospora crassa* EAS; *Cryphonectria parasitica* CRP (Cryparin); *Ophiostoma ulmi* CU (Cerato ulmin). Photo copyrights: Kimberlie Sasan (top) Matt Kasson (bottom). Links and additional information for the images including their associated CC licenses are provided in Supplemental File 1.

Other proteins would later join these *S. commune* hydrophobins to establish the hydrophobin family (**Figure 1 B-E**). At the time, a wide range of small, secreted proteins predicted from mRNAs were identified, all containing an 8-Cys pattern with conserved relative spacing. These were abundantly expressed at various stages of development in filamentous fungi across Dikarya (Ascomycota and Basidiomycota). It is interesting to note that previously experimentally characterized proteins would join the hydrophobin family at this time on the basis of their 8-Cys pattern. These included the *Ophiostomo ulmi* virulence factor cerato ulmin (CU) (Stringer and Timberlake, 1993), the host cell-wall associated *C. parasitica* cryparin (CRP)(Zhang et al., 1994), and the spore-associated *Neurospora crassa* EAS (Bell-Pedersen et al., 1992)(Lauter et al., 1992).

Together, these sequences established our view of the hydrophobin family and their canonical definition in use today (**Box 1**). In addition to their conserved 8-Cys pattern, the hydrophobin sequences contain an N-terminal secretion signal that is cleaved upon export into the extracellular space. Considering their mature sequences (i.e., without secretion signal), there is limited sequence conservation across the family. For instance, EAS and RodA share as little as 10% identity in their alignments despite belonging to the same protein family. On the other hand, certain sequences, such as CRP and CU share more sequence identity (48%) and identical cysteine residue spacing. This disparity in sequence conservation led to the subdivision of the hydrophobin family into two Classes, I and II. CU and CRP would be assigned the Class II subdivision while the remaining highly diverse sequences would be assigned to Class I. Later studies would ascribe distinct properties to the assemblies formed by members of the two classes: Class I hydrophobins form exceptionally stable amyloid-like rod-shaped structures that can only be disassociated with strong acids, while Class II hydrophobins form (comparatively) less stable assemblies that can be disassociated with alcohol-detergent mixtures (**Box 1**).

#### Box 1

##### Canonical Definition for the Hydrophobin Family and its Subdivisions (Wessels et al.,1997)

###### Hydrophobin protein sequence

Moderately hydrophobic small secreted proteins (~ 100 amino acids) with a strictly conserved eight-cysteine pattern which otherwise, considered together, have poor amino acid homology. There is conservation, however, in their hydropathy profiles (i.e. the relative position of regions with overall more hydrophobic or hydrophilic amino acids) when the strictly conserved cysteines are aligned (**Figure 1**).

###### Hydrophobin behavior in solution

When presented with a hydrophobic-hydrophilic interface, such as the air-water interface, or a surface, hydrophobins self-assemble to form amphipathic films i.e. one side is more hydrophobic than the other.

**The canonical hydrophobin family can be subdivided into two classes (I and II)**.

###### Class I hydrophobins

Form highly insoluble rod-shaped nanostructures (rodlets) with amyloid-like structure (Mackay et al., 2001) that can only be dissociated by strong acids such as formic acid.The spacing between the cysteines is highly variable.

###### Class II hydrophobins

Form molecular films that can be disrupted by alcohol-detergent mixtures (do not have rodlet morphology). The spacing between the cysteines is highly conserved.

While the foundational hydrophobin amino acid sequences are highly diverse, the relative position of the hydrophobic and hydrophilic amino acids within the amino acid sequence (hydropathy plots) are similar. Initially, the comparison of hydropathy plots was used to identify regions that potentially share similar three-dimensional structures within the sequence (i.e., hydrophobic amino acids are more likely to be buried within the protein core than hydrophilic residues which tend to lie at the surface of the protein (Olsen, 1980)).

More recently, the molecular structures of a number of diverse hydrophobin sequences have been experimentally determined by crystallography and/or solution NMR (Gandier et al., 2017; Hakanpää et al., 2006, 2004; Morris et al., 2013; Pham et al., 2016; Ren et al., 2014; Valsecchi et al., 2020), revealing a shared generic structure: a small β-sheet core, whose curvature can vary, with highly variable peripheral secondary structures (**Figure 2**). Interestingly, the hydropathy pattern previously used to describe the hydrophobin family is a reflection of the universally conserved four β-strands that assemble together to form the β-sheet core. Furthermore, it was found that the 8-Cys pattern forms four strictly conserved disulfide bridges. Two of the four disulfide bridges form between consecutive β-strands likely stabilizing the protein’s core. The remaining two disulfide bridges form between the protein’s β-sheet core and the “loops” formed by the intervening regions between the β-strands. These modular loop regions accommodate the remarkable sequence diversity displayed by the family, as a wide range of structures or dynamic loops can be accommodated within these spaces (Gandier et al., 2017; Pham et al., 2018). In class II hydrophobins, these regions are shorter than those in class I, explaining in part the better quality of their sequence alignments. Furthermore, the class II structure is globular and compact while class I structures can be seen as more dynamic in solution. These features are also reflected in their distinct molecular assembly mechanisms. Class II assembly as it is understood is driven by a hydrophobic patch at the protein’s surface (Paananen et al., 2021; Szilvay et al., 2007). Such a patch has yet to be discovered at the surface of a Class I hydrophobin. While the mechanism by which Class I hydrophobins functionally assemble have not been fully elucidated, evidence is accumulating that significant structural shifts contribute to the process (Macindoe et al., 2012; Valsecchi et al., 2020).

**Figure 2.**
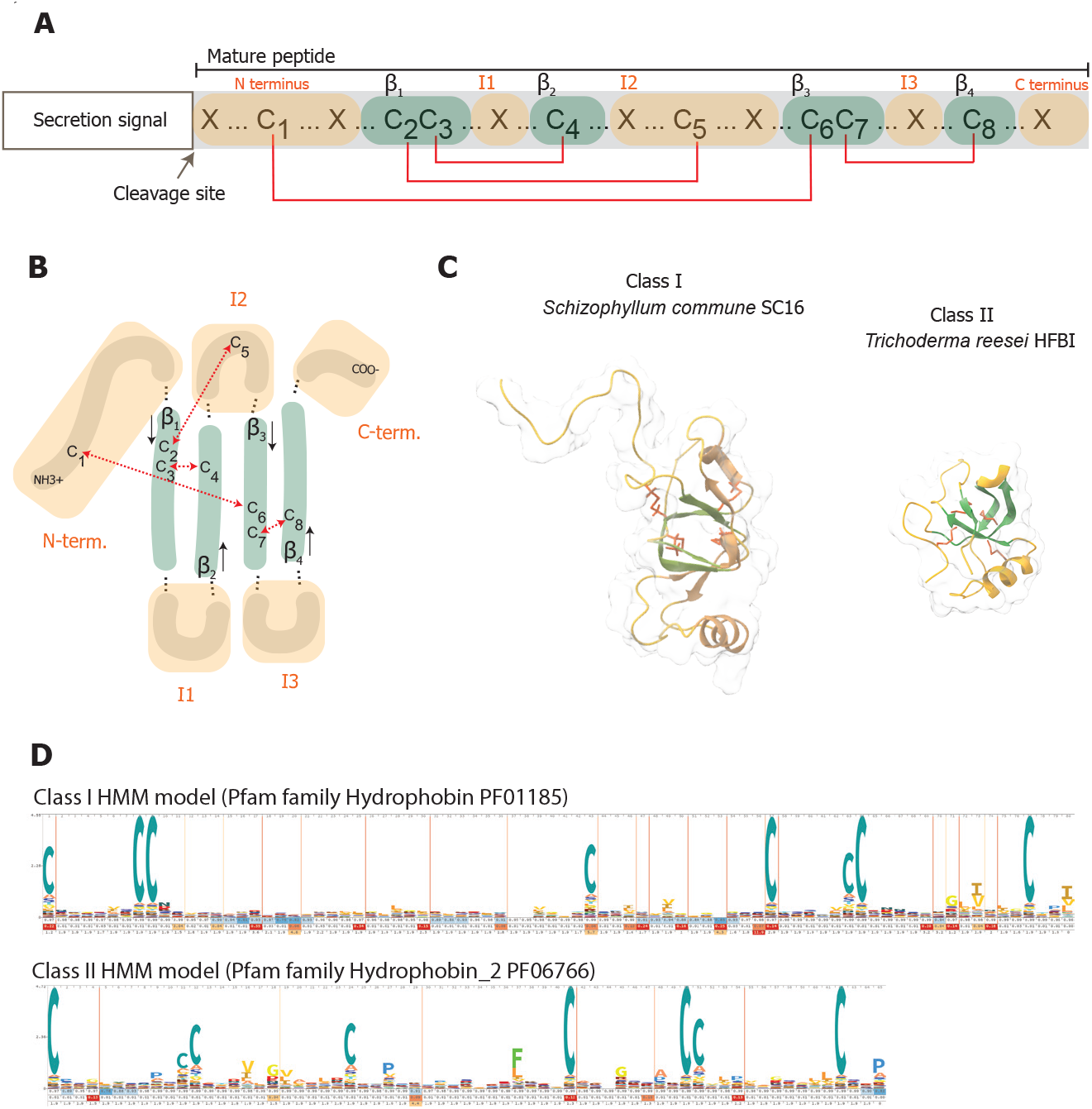
The generic structure of the hydrophobin family and examples of Class I and II molecular structures. (A) Hydrophobins are low molecular weight proteins characterized by a strictly conserved spacing of eight cysteine residues that form four disulfide bridges (indicated by red brackets). (B) All hydrophobin structures that have been experimentally resolved can be described by a generic two-dimensional topology centered on four β-strands that organise into an anti-parallel sheet indicated in green. The intervening regions between the β-strands (I1, I2, and I3) as well as the N- and C-termini are highly variable in their chemistries and structures (or lack thereof) and are indicated in orange. The four conserved disulfide bridges within the 8-Cys pattern are indicated by red arrows. (C) Simplified ribbon backbone representations of two experimentally determined Class I and II molecular structures colored to correspond to the generic structure: *Schizophyllum commune* Class I hydrophobin SC16 (Gandier et al., 2017) (RCSB PDB DOI: 10.2210/pdb2NBH/pdb) and Class II *Trichoderma reesei* hydrophobin HFBI (Hakanpää et al., 2006) (RCSB PDB DOI: 10.2210/pdb2FZ6/pdb). (D) Logos for the HMM Pfam models for Class I (PF01185) and Class II (PF06766). The unifying feature is the 8-Cys pattern. Intervening regions and chemistry are comparatively more conserved in Class II sequences than Class I sequences. This can be seen in the limited conservation displayed by the class I sequence model logo.

## 4. Methods

### 4.1 Proteome selection

A total of 52 proteomes were included in this analysis from a range of organisms available from multiple sources (i.e. JGI, NCBI, and our ongoing sequencing projects). Proteomes were chosen to span the breadth of the Kingdom Fungi and other relevant organisms that co-occur with fungi across multiple environments and variously represented fungus-like lifestyles (i.e., sporulation and filamentous growth). This includes Oomycetes (water molds), Amoebozoa (slime molds), Actinobacteria (filamentous bacteria), Apicomplexa (*Plasmodium*), and Archaea (methanogens). These proteomes and their metadata, including accession numbers and source, are reported in **Supp. File 1**. The 45 representative fungal genomes span eight phyla and more than 25 orders, but nearly 40% are contained within two fungal classes: the Agaricomycetes (Basidiomycota) and the Sordariomycetes (Ascomycota). Approximately 15% of all fungal genomes included in this study are members of the order Hypocreales (Sordariomycetes) due in part to their diverse ecologies and importance as disease-causing agents in both plants and animals. The various ecologies of the surveyed fungi spanned numerous plant and animal pathogens and mutualists, mycoparasites, decay fungi, and edible fungi (**Figure 3**). Each organism was assigned a three-letter acronym consisting, when possible, of the first letter of the genus followed by the first two letters of the species (**Supp. File 1**). The acronyms for organisms included in this hydrophobin survey are provided in **Table 2**.

**Table 2.**
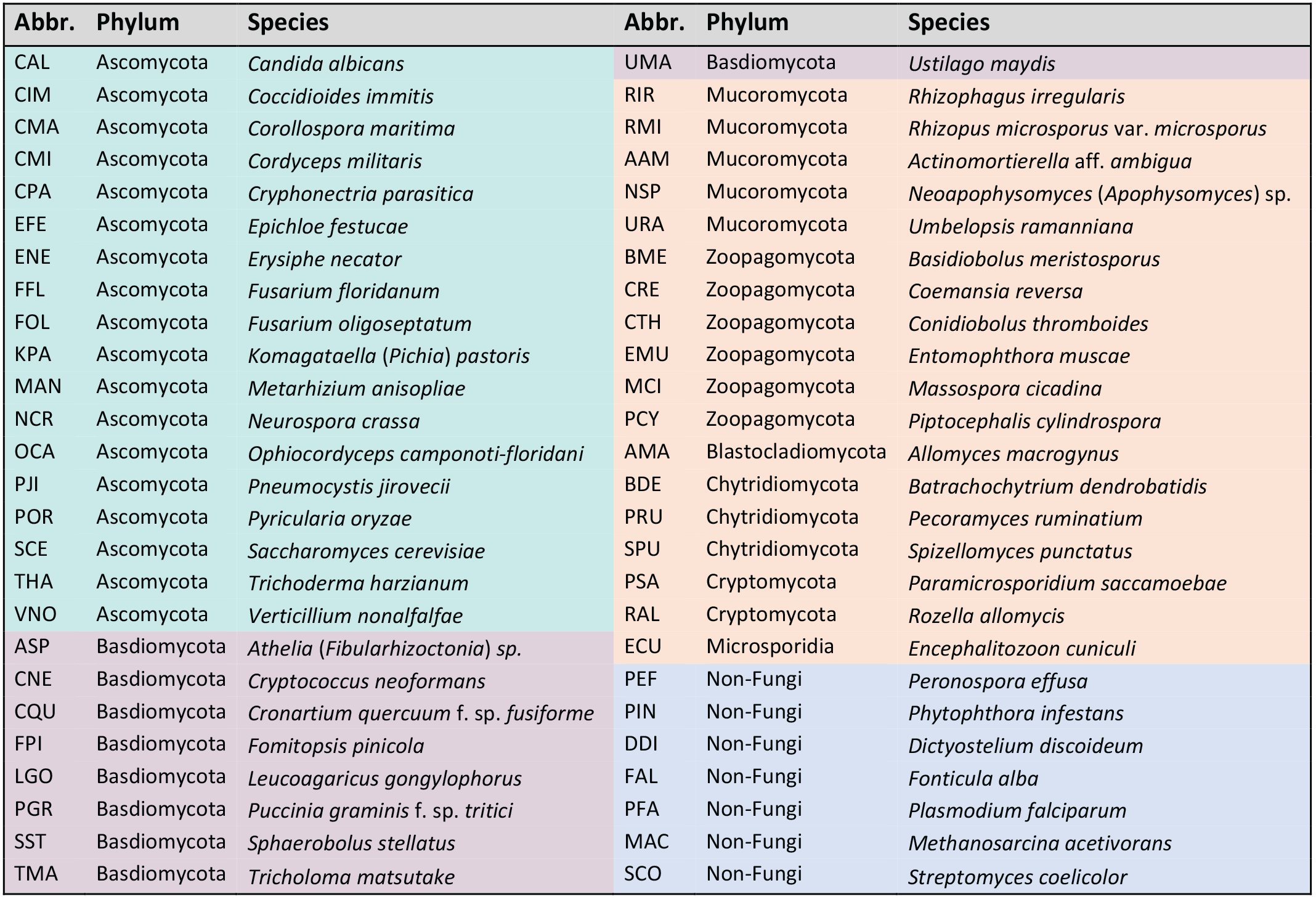
A complete list of organisms included in this study.

**Figure 3.**
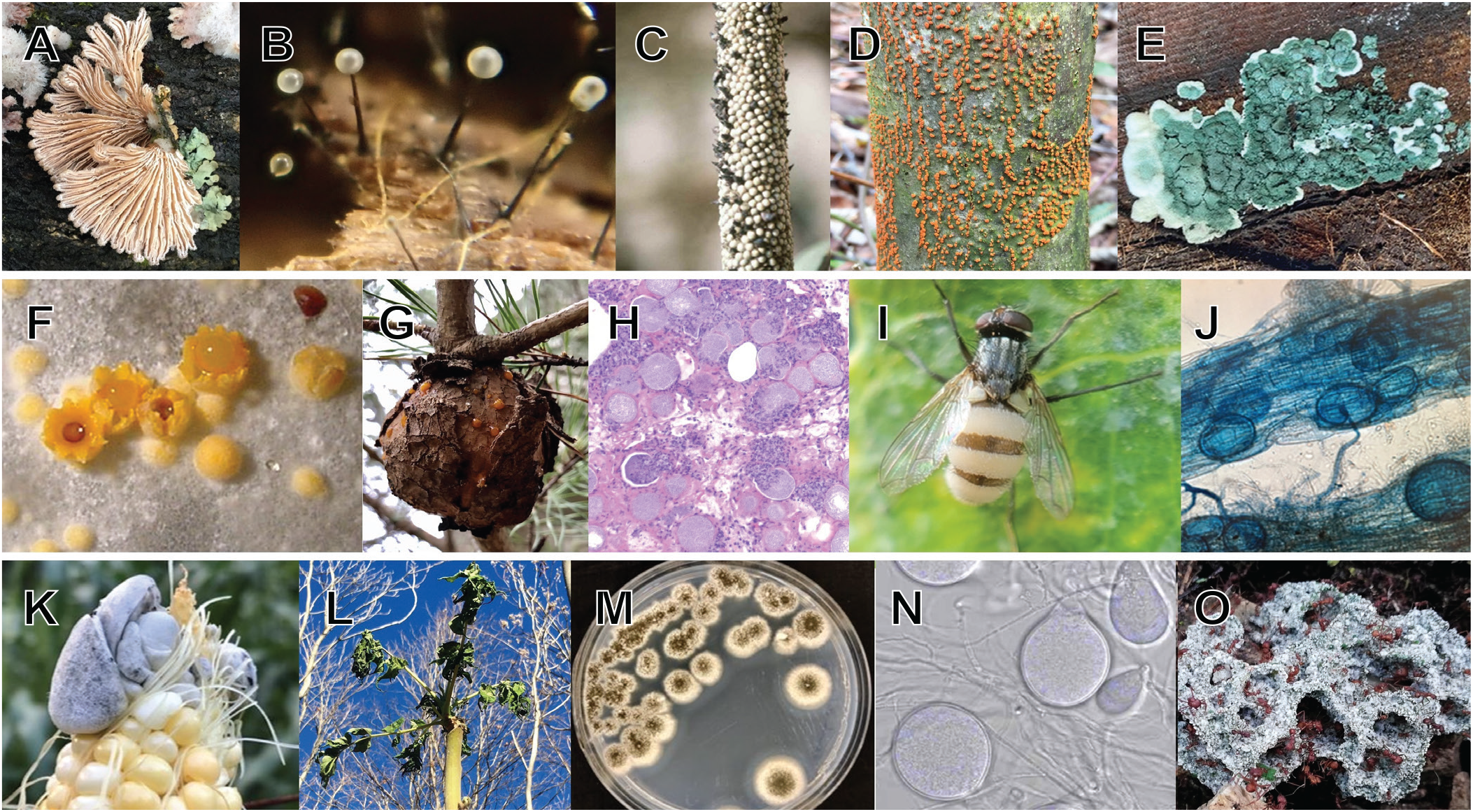
Representative Fungi included in Kingdom-wide survey for hydrophobins and hydrophobin-like proteins. A-E these fungi contain well characterized hydrophobins that served as positive controls/indicator sequences in our study; C-O fungi included in our survey. Fungal identifications are as follows: A. *Schizophyllum commune*, B. *Ophiostoma novo-ulmi*, C. *Claviceps fusiformis*, D. *Cryphonectria parasitica* E. *Trichoderma sp*., F. *Sphaerobolus stellatus*, G. *Cronartium quercuum*, H. *Coccidioides immitis*\*, I. *Entomophthora muscae*, J. *Rhizophagus irregularis*\*, K. *Ustilago maydis*, L. *Verticillium nonalfalfae*, M. *Metarhizium anisopliae*, N. *Pecoramyces ruminatium*, O. *Leucoagaricus gongylophorus*; Photos A-C, E, H, K, and N-O were cropped / rotated to enhance certain features. Photos denoted with an asterisk indicate closely related species of the same genus are shown in photos. Photo copyrights: A. John Boback, B. Nur Ritter, C. USDA ARS, D,G, J, L, and M Matt Kasson; E, lilithohlson, F, Jozsef Geml, H. Bridget Barker, I. Gene H., K. Brian Hunt, N., Noha Youssef, and O. Lizette Cicero. Links and additional information for the images including their associated CC licenses are provided in **Supp. File 1**.

### 4.2 Identification of experimentally confirmed hydrophobin sequences (indicator sequences)

Given that the annotations of protein sequences in genomic and proteomic databases are based primarily on bioinformatic predictions, the literature was surveyed to identify experimentally confirmed protein sequences that could serve as reference points to both help identify and contextualize novel hydrophobin sequences in our downstream methodologies. It is important to note that the primary criteria for these sequences was the existence of experimental evidence for their biophysical properties. While this approach limits the scope of the indicator sequences to those whose properties have been directly observed (e.g., within Dikarya), it also provides higher confidence in terms of hydrophobin sequence candidacy. Of the total indicator sequences described in **Table 1**, 22 were chosen as they have direct experimental evidence for the properties of their protein assemblies such as their solubility, thus distinguishing their canonical classification (Class I or II). Two were chosen since their molecular structures have been experimentally determined and are consistent with the generic hydrophobin structure, as well as Class specific features (these are defined in **Figure 2**). Finally, an additional five indicator sequences were included (RodC-G) to complete the suite of hydrophobins encoded in the *Aspergillus fumigatus* genome whose hydrophobin knock-outs have recently been systematically studied. Together, a total of 29 indicator sequences were included. A detailed list of their sequence names, source fungus, molecular characteristics as well as biological characteristics are described in **Table 1**. All indicator sequences included in this study are known to be secreted and so were trimmed for downstream analysis based on SignalP (ver. 5.0 for Eukarya) (Almagro Armenteros et al., 2019). When possible, the site of cleavage upon secretion was cross-referenced with experimentally determined mature protein sequences. It is important to note that the selection of organisms whose proteomes are included in the study was made independently of the selection of indicator sequences. Of the indicator sequences, four are found in proteomes that are also included in the study (i.e., CPA, POR, NCR, and MAN).

### 4.3 Permissive mining of selected proteomes for the construction of a working database of protein sequences with features associated with hydrophobins

The proteomes of the selected organisms were subjected to a custom bioinformatic pipeline summarized in **Figure 4**. First, the proteomes were permissively mined to build a database of sequences with features broadly associated with hydrophobins (**Figure 4A-C**). To achieve this, a search exclusively based on the presence and relative spacing of the central motif of the 8-Cys pattern (Cys2 to Cys7) was conducted. This motif is referred to hereafter as the 6-Cys pattern (C2C3 -X- C4 -X - C5 -X- C6C7; where X represents 1 or more of any amino acid, except Cys). No constraints were imposed on the distance between cysteine residues, maximal number of cysteines, or protein length. A custom R function relying on the package effectR (Tabima and Grünwald, 2019) was used to pull these candidates. To ensure sequences following the canonical definition of the hydrophobin sequence were also captured in the database, hmmsearch (ver. 3.1b2) with the permissive default e-value threshold of 10 was used to pull sequences annotated as hydrophobins using Pfam models (ver. 34.0) of Class I (PF01185) and Class II (PF06766). (Tabima and Grünwald, 2019)The combined results of these searches were then submitted to SignalP (ver. 5.0 for Eukarya) (Almagro Armenteros et al., 2019) to identify secretion signals. When present, these were trimmed such that only the mature protein sequences are included in the study. These were then additionally annotated with hmmsearch against Pfam-A models (ver. 34.0). Finally, the experimentally validated, mature indicator sequences (**Table 1**) were combined with the mined sequences. Together, these form the working database of sequences.

**Figure 4.**
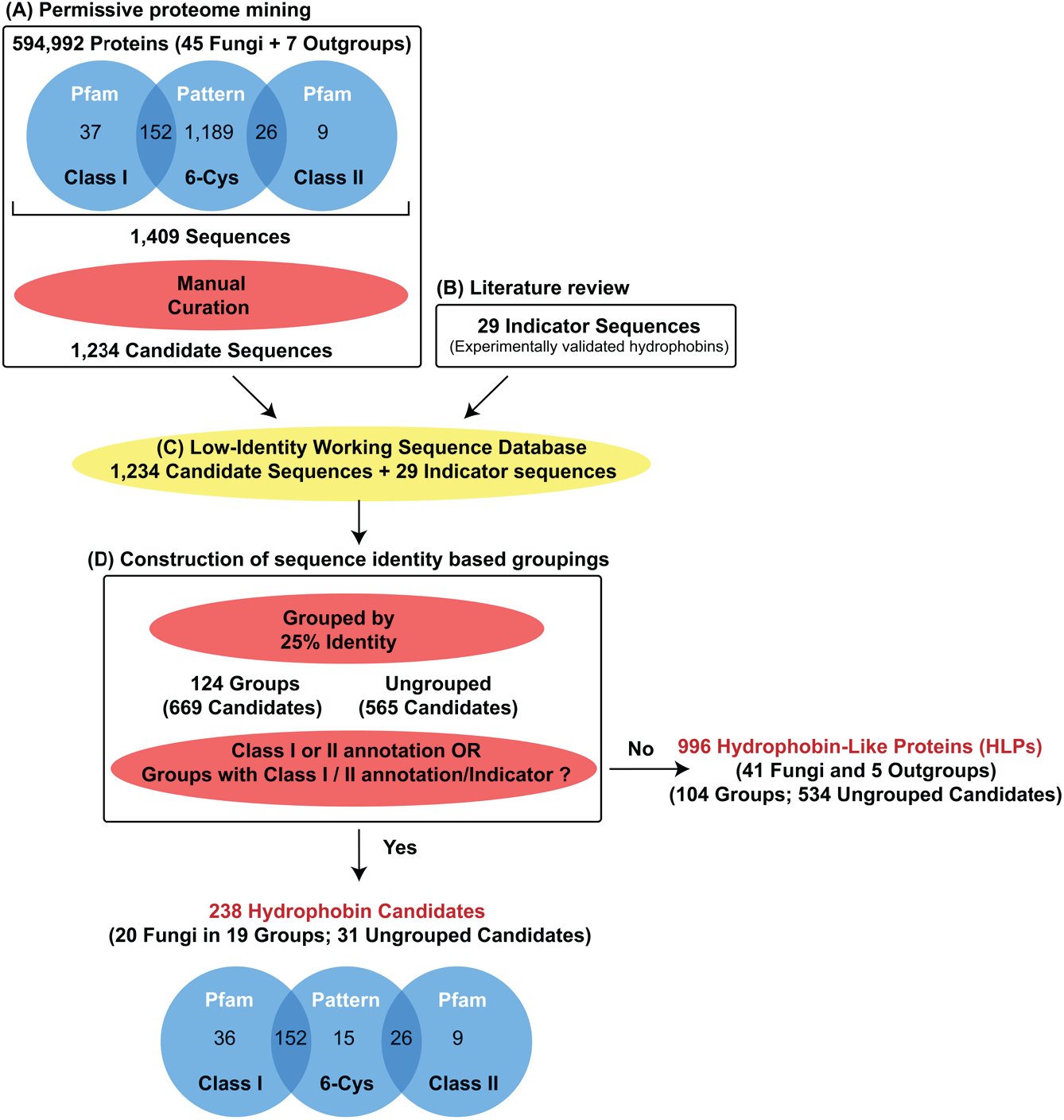
Overview of the workflow used to establish the cross-Kingdom hydrophobin candidate identity groupings. (A) The proteomes of 52 organisms from within and outside Kingdom Fungi were permissively mined for hydrophobin-like features by searching for a truncated form of the 8-Cys pattern referred to herein as the 6-Cys pattern. These were manually curated to remove excessively cysteine-rich proteins as well as those sequences whose cysteines participate in distinct patterns known to belong to other well-established protein families. Proteomes were also submitted to Pfam for sequence-based prediction of canonical Class I (PF01185) and Class II (PF06766) domains. (B) These sequences were combined with the sequences of experimentally validated hydrophobins (indicator sequences) identified in the literature to build (C) a database of sequences with a broad range of amino acid chemistries and structures. (D) The sequences in the database were grouped together with a threshold of 25% sequence identity. Groups that contained an indicator sequence and/or Pfam hydrophobin annotation were defined as being composed of hydrophobin candidates. Ungrouped sequences annotated as hydrophobins by Pfam were also considered hydrophobin candidates. The remaining sequences were defined as hydrophobin-like proteins (HLPs).

### 4.4 Construction of sequence identity-based groupings distinguishing hydrophobins candidates from hydrophobin-like proteins (HLPs)

To understand how the sequences contained within the working database might relate to each other, sequences were grouped according to a sequence identity threshold (25%) using a sequence similarity network approach (Copp et al., 2018)(**Figure 4D**). To do so, a MAFFT alignment (ver. 7.450) (Katoh and Standley, 2013) of the working database was produced, then custom R scripts (ver. 4.1.0) (R Core Team, 2021; RStudio Team, 2021) were used to identify then group together all sequences that share at least 25% identity. This low identity threshold captured observable clusters of identity in full alignment matrix and also allowed clustering of at least 50% of sequences. Through this approach, criteria were established to identify sequences within the working database that can be associated with the canonical hydrophobin definition. These are referred to as hydrophobin candidates and were defined as having at least one of the following features: groups with an indicator sequence, groups with a sequence containing a hydrophobin Pfam annotation, is an individual sequence that contains a Pfam annotation itself. Sequences that do not meet the criteria outlined in this definition are referred to as hydrophobin-like proteins (HLPs). In addition, using the built-in R function, principal component analysis was performed on the hydrophobin candidate subset of the pairwise sequence identify matrix generated from the MAFFT alignment.

### 4.5 Construction of strict hydrophobin group HMM profile library and pairwise co-classification

For each hydrophobin candidate group, a MAFFT alignment was constructed and used to generate Hidden Markov Model (HMM) profiles along with their related sequence logos using Skylign (Wheeler et al., 2014) (removing mostly empty columns and with above-background letter height). The workflow of subsequent steps is described in Figure 5. The strict group HMM profiles along with the existing Class I and Class II Pfam models (PF01185 and PF06766, respectively) were combined into an HMM library. This profile library was applied using hmmsearch with default parameters (e-value threshold of 10) to re-assign the HMM-defined groups for hydrophobin candidate sequences. This step was taken to identify which hydrophobin candidate sequences could be classified within multiple HMM-assigned groups and whether the models contained enough sequence information to reproduce the original groupings. An R script was written to rank the scores generated by each model for each sequence. We refer to the relationship between a single sequence that can be captured by multiple models in the HMM library as co-classification. We further quantified overlap between groups by calculating the percent overlap of shared sequences in two groups. Specifically, for each group, the number of proteins in one group that co-classified into another group was divided by the total number of proteins within the initial group. All scripts and files generated for this analysis are available at an archived repository for this project (Lovett, 2022).

**Figure 5.**
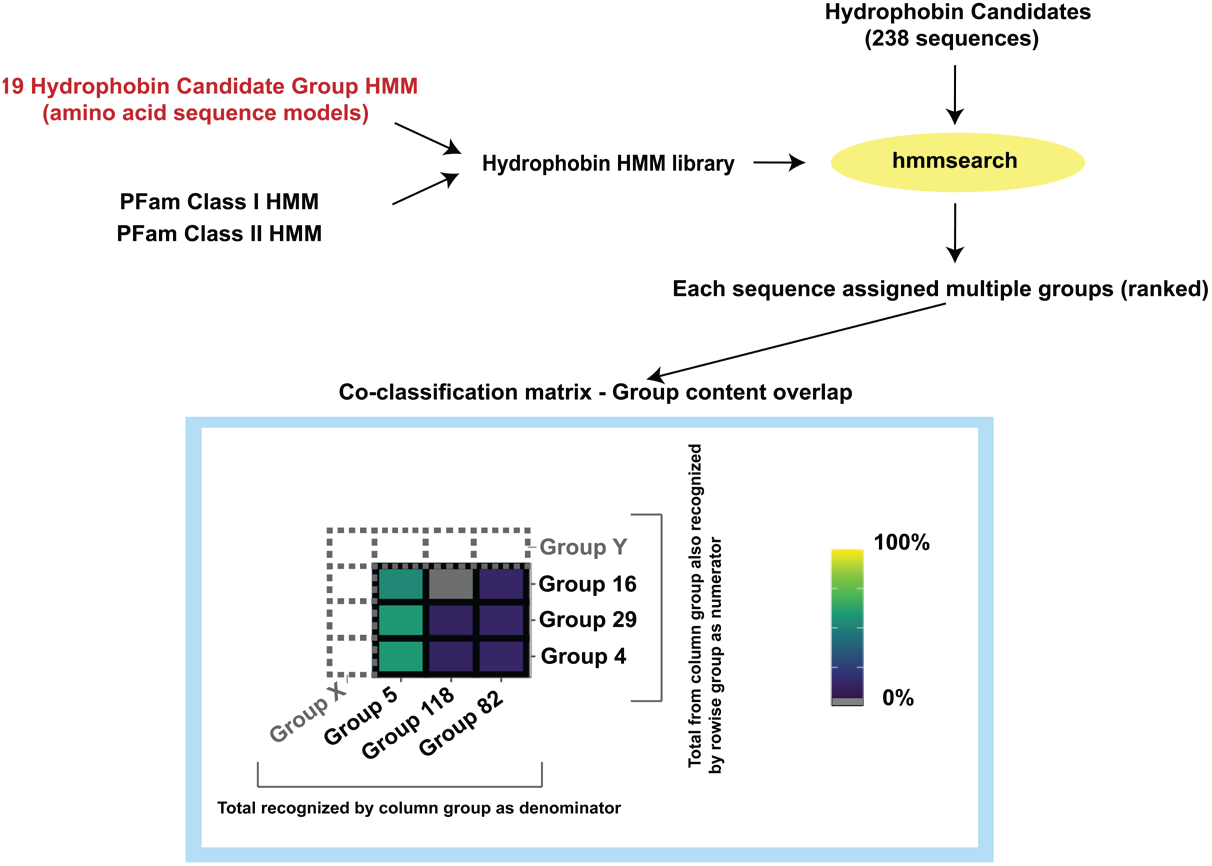
Overview of the workflow used to establish and study the library of overlapping hydrophobin candidate HMM profiles. For each group previously identified as consisting of hydrophobin candidates (**Figure 4**) an HMM profile was generated from its sequences. These profiles were combined with the Class I and II Pfam profiles to generate the hydrophobin HMM profile library. This library was then used by hmmsearch to assign sequences to group profiles and/or Pfam classes (e-value threshold of 10). The co-classification matrix of all proteins within each group was then constructed as a means of visualising the network generated by the overlapping classifications of sequences within groups. Values (% overlap) are reported according to the column (i.e., percent of members from the columnar group that were also present in the row-wise group). Gray boxes signify no overlap in group content.

## 5. Results

### 5.1 A working database of sequences was constructed for the study representing a highly diverse subset of predicted proteins with features similar to those of hydrophobins

The permissive screen of proteomes for hydrophobin-like features allowed the construction of a working database of 1,234 highly diverse sequences within and beyond Kingdom Fungi (**Figure 4A-C**). More specifically, the working database consists of sequences pulled from all 45 fungi and 7 outgroup organisms included in the study, as well as 29 experimentally characterized indicator sequences summarized in **Table 1**. It is important to note that the aim of this approach was not to be exhaustive, but rather to build a dataset that begins to explore the potential range of amino acids that can be accommodated in the modular structure of the hydrophobin sequence. In fact, it was anticipated during the design of the custom bioinformatic pipeline that the working database would not exclusively consist of hydrophobin candidate sequences, nor be necessarily related to hydrophobin sequences beyond the search criteria that define their inclusion (e.g., 6-Cys motif with no other constraints). In other words, the aim was to create a subset of highly diverse cysteine-rich proteins to study. This is reflected in the MAFFT alignment of the sequence database (pairs of compared sequences ranging from 0-100% identity) and subsequent construction of the sequence identity-based groupings (referred to hereafter as groups) (**Figure 4D**). An identity threshold of 25% grouped 54% of the sequences contained in the working database, the remaining 46% of the sequences could not and remained ungrouped.

Of the total sequences in the working database, 238 were determined to be hydrophobin candidates. To qualify as such, a given sequence must share a group with an indicator sequence, share a group with a sequence that is annotated by Pfam as a hydrophobin, or is annotated itself by Pfam as a hydrophobin. The distribution of core hydrophobin candidates by organism and by the screening criteria that initially pulled their sequences into the working database are described in **Figure 6**. There was no overlap between sequences pulled into the working database by Pfam Class I annotations and Pfam Class II annotations. There was, however, near-entire overlap between the sequences pulled by the presence of the 6-Cys motif and Pfam annotations. While only 15 hydrophobin candidates were pulled exclusively by the permissive 6-Cys motif, it is important to note that these sequences would otherwise not have been associated with the hydrophobin family. The remaining working database sequences, a total of 996 excluded by the conservative hydrophobin candidate definition, are referred to as hydrophobin-like proteins (HLPs). The extent to which HLPs may or may not be related to the hydrophobin family was not investigated in this study.

**Figure 6.**
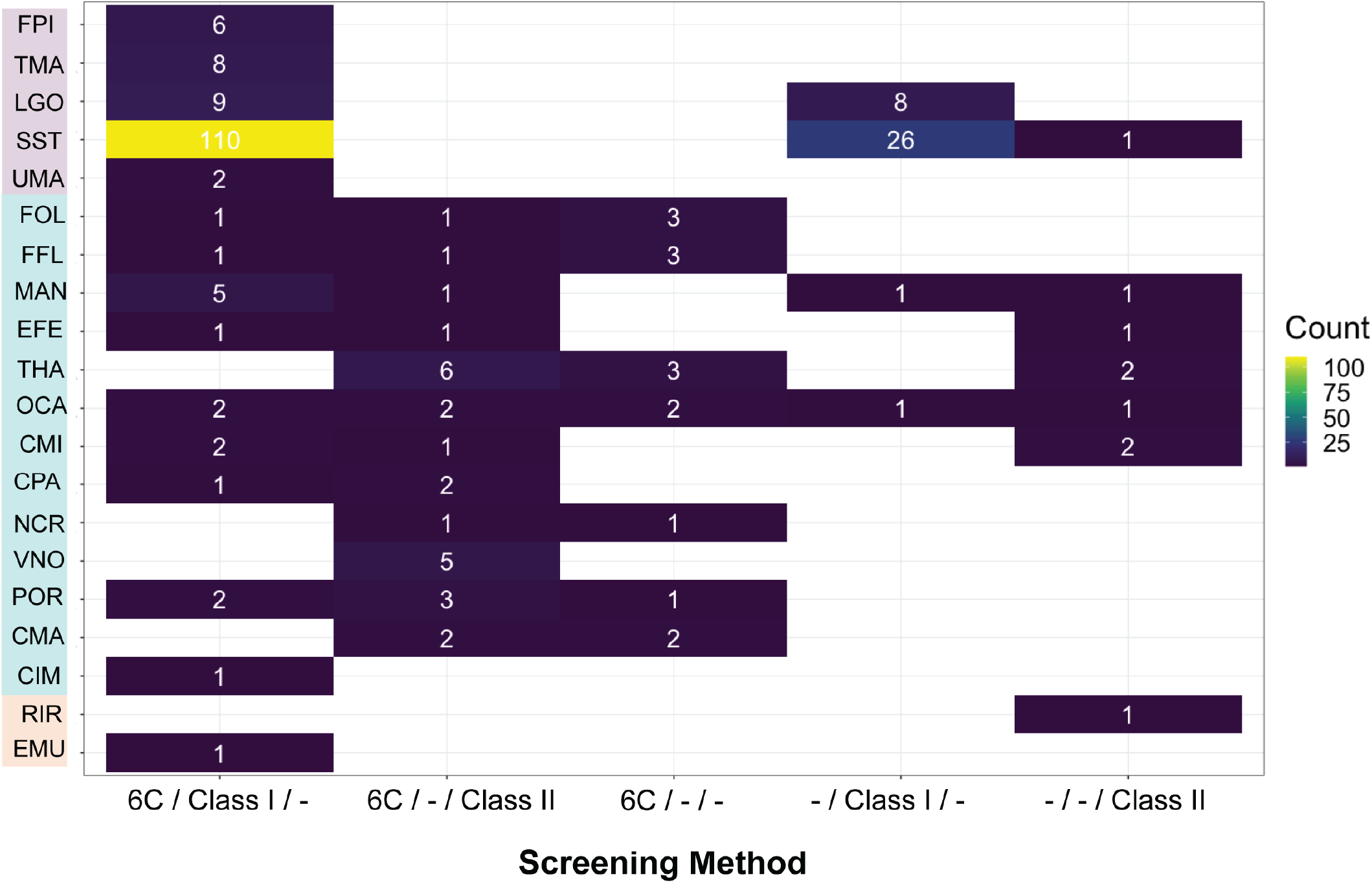
Distribution of the groups consisting of hydrophobin candidates amongst the organisms included in the study and by screening method. Only the organisms from which hydrophobin sequences were pulled are listed (full names listed in Table 2). Three methods were used to screen the proteomes included in the study: the permissive 6-Cys pattern, class I Pfam HMM model, class II Pfam HMM model. A sequence could be pulled by both a Pfam model and 6-Cys pattern search. There was no overlap between search results using Pfam models for Class I and II.

It is important to note that we rely on sequence identity (i.e., count of matching sequences) rather than similarity (i.e., calculated via sequence distance) to build these groups of sequences, as our ultimate aim is to describe the full spectrum of potential hydrophobin amino acid sequence chemistries within the working database. We did not want to build-into this analysis assumptions of similarity built into existing similarity scoring systems.

### 5.2 The inclusion of experimentally validated canonical Class I and Class II hydrophobin indicator sequences in the working database provided context for the sequences with which they form groups

The nature of the groups formed in the study could be assessed within the context of the established hydrophobin literature, as the experimentally validated indicator sequences were included in the alignment and appeared in the subsequent groupings. The groups for each indicator sequence along with their experimentally validated properties are reported in Table 1. Overall, the groups reflected the canonical class dichotomy: Class I indicator sequences did not form groups with Class II indicator sequences. Similarly, no group contained sequences with both hydrophobin Pfam annotations. It is important to note that canonical classes were assigned to each hydrophobin candidate group based on these existing annotations and/or the presence of an experimentally validated indicator sequence.

Given their conserved nature, the Class II indicator sequences grouped exclusively together as anticipated (group 69: CFU_TH1, CPA_CRP, MOR_MHP1, NCR_NC2, TRE_HFBI, TRE_HFBII + 21 candidate sequences), with the exception of *O. ulmi* cerato-ulmin (OUL_CU). Instead, OUL_CU formed the separate Group 1 with a single candidate *T. harzianum* (XP_024771806-1). In contrast to the Class II indicator sequences, Class I indicators are found across a wide range of groups. Of these, Group 10 contained the highest number of Class I indicators with a total of four (132 candidates, POS_VMH2, SCOM_SC3, SCOM_SC4 and GFR_HGF1).

Three groups contained two indicator sequences: Groups 117 (6 candidates, SCOM_SC16, MAN_SSG1), Group 79 (2 candidates, XPA_XPH1, MOR_MPG1), and Group 118 (1 candidate; AFU_RodC, AFU_ RodA). The latter Group 118 contains a single candidate from *Coccidioides immitis* (CIM_XP_001242719). While groups containing experimentally confirmed Class I indicators also contained sequences with Class I PFam assignations, Group 5 did not. This group consisted of the Class I indicator TAS_TasHyd1 along with six candidates that were pulled into the analysis using the permissive 6-Cys pattern screen. These strict candidates represent five Sordariomycetes genomes, including one from the marine fungus *Corollospora maritima* and two candidates from *Trichoderma harzianum*. Group 85 consisted of the indicator (PCA_ HYDX) and a single candidate from *Sphaerobolus stellatus*.

The remaining indicators included in the study did not group with other candidate sequences (ANI_DewA, AFU_RodD, AFU_RodG, AFU_RodB, AFU_RodF, SLA_HYDX, WIC_HYDX). The sequences AFU_ RodE and NCR_EAS grouped exclusively with homologs (groups 39 and 82, respectively). NRC_EAS was not captured by the Pfam HMMs for hydrophobins, as permissively applied in the study, although it is one of the most rigorously experimentally characterized hydrophobins (**Table 1**).

### 5.3 Hydrophobin candidate sequences were limited to fungi, but identified within and beyond Dikarya

Some hydrophobin candidate groups have clear phylogenetic signals. The 238 hydrophobin candidates recovered formed a total of 19 groups (non-consecutively numbered) whose distribution across 20 fungal proteomes is described in Figure 7. Though mostly found in either Ascomycota or Basidiomycota, these hydrophobin candidates were also present outside of these phyla, namely the Mucoromycota and Zoopagomycota. The Ascomycota surveyed yielded the highest percent incidence of strict hydrophobins with 72% (13 out of 18 genomes), followed by Basidomycota 55% (5 of 9 genomes), Mucoromycota 20% (1 of 5 genomes), and Zoopagomycota 17% (1 of 6 genomes). The four other fungal phyla (Cryptomycota, Blastocladiomycota, Chytridiomycota and Microsporidia) did not yield hydrophobin candidate sequences, nor did any of our chosen outgroups (Oomycota, Apicomplexa, archaea, bacteria and Amoebozoa).

**Figure 7.**
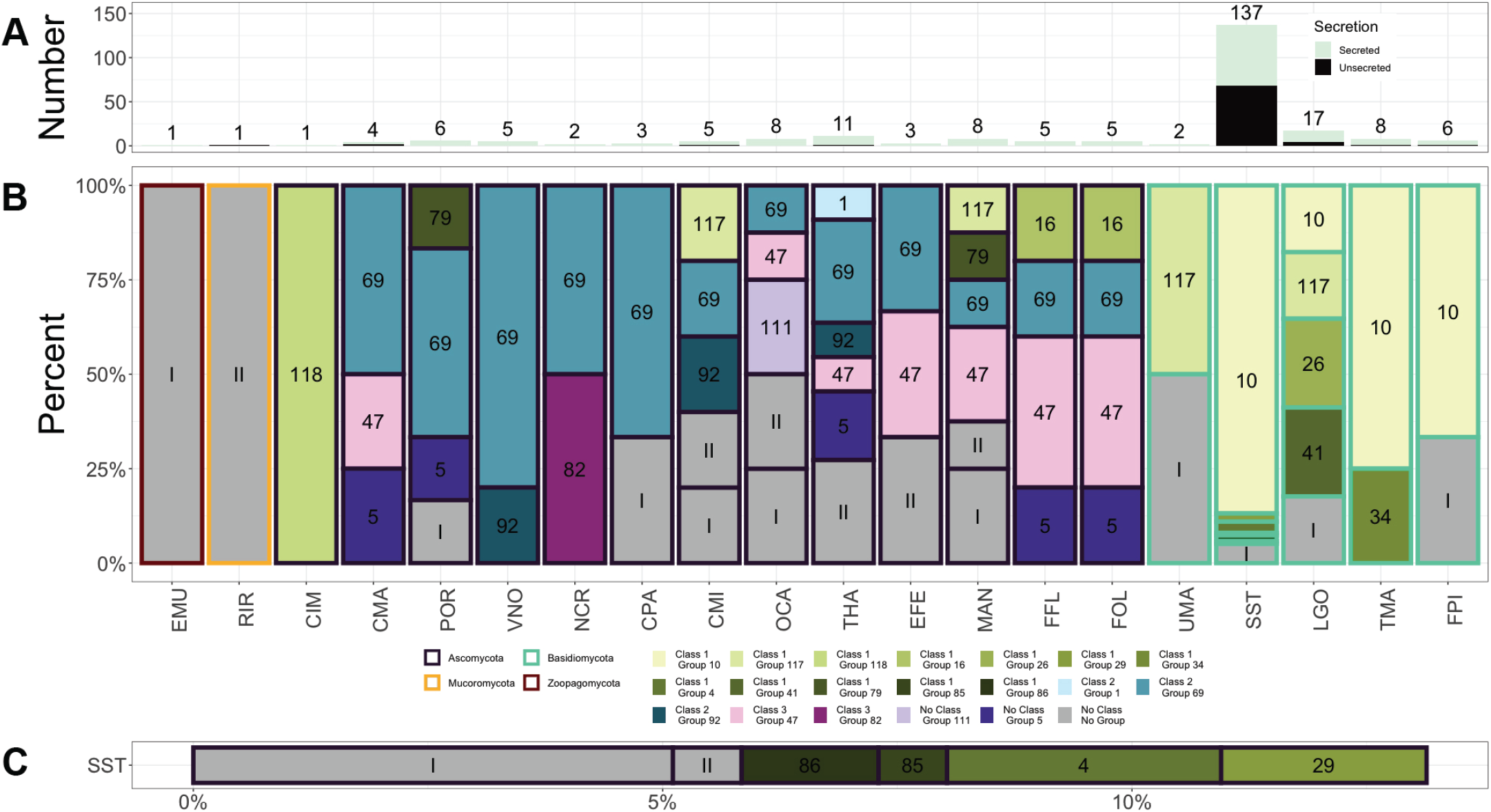
Ascomycota and Basidiomycota have distinct hydrophobin grouping profiles containing sequences with and without predicted secretion signals. (A) Overall abundance of hydrophobin candidates across Kingdom Fungi. Proportion of sequences with predicted secretion signals indicated by relative proportion of light green colour in respective bar. (B) Distribution of hydrophobin groups across fungi divided by color based on the class ontology developed in this study (Section 5.9; Class 1 in green; Class 2 in blue and Class 3 in red). Groups within classes are distinguished by shade. Unclassified and ungrouped candidates are marked in grey with roman numerals indicating Pfam classes. (C) Distribution of SST groups, excluding Group 10, which was notably inflated in this fungus.

It is interesting to note that compared to other Dikarya, a larger proportion of hydrophobin candidates from Sordariomycetes (Ascomycota), did not form groups.

Overall, the survey shows clear evidence that hydrophobins expand beyond the bounds of Dikarya, including one representative each from Zoopagomycota (EMU; an entomopathogenic fungus) and Mucoromycota (RIR; an arbuscular mycorrhizal fungus). The 29 ungrouped Dikarya hydrophobin candidates were evenly distributed among Basidiomycota and Ascomycota. In contrast to grouped hydrophobins, a disproportionate number of ungrouped hydrophobins (~25%) did not contain a predicted secretion signal. Such ungrouped intracellular hydrophobins were found from one fungus outside of Dikarya (RIR), two representatives from Basidiomycota (SST and LGO), and one representative from Ascomycota (CMI).

Of the hydrophobin candidate groups, eight contained sequences exclusive to Basidiomycota, while ten were exclusive to Ascomycota. A single group (117) spanned genomes from both Basidiomycota (LGO) and Ascomycota (CMI and MAN). This group represented the only hydrophobin candidate group maintained across Dikarya. This finding is underscored by the two indicator sequences assigned to Group 117 originating from fungi in these different phyla (i.e., *Schizophyllum commune* and *Metarhizium anisopliae*). Only two Basidiomycota groups spanned multiple genomes (Group 10 and Group 117), compared to seven Ascomycota groups. Nearly all nineteen groups contained at least one sequence with a secretion signal. Group 4 (exclusive to SST), was composed of four sequences without predicted secretion. Five other groups had at least one accession lacking a secretion signal: four of these were exclusive to Basidiomycota. Among strict hydrophobins, 31 (~13%) did not group with other sequences.

A number of the groups established in the study showed clear phylogenetic signals. Group 10, for example, was limited to, but widespread within, Basidiomycota. Groups 69 was present in all Ascomycota that contained hydrophobin candidates, except the dimorphic human pathogen CIM. Group 47 was present across all hypocrealean fungi, but also CMA, a cosmopolitan marine fungus in the Microascales. Group 5 was found from only, but not all, plant-associated fungi, including CMA which is found in living tissues of seagrass (Cuomo et al., 1985) to ultimately degrade plant material as a saprotroph (Raghukumar, 2017) (Li et al., 2016). The two closest related species in our analysis were ambrosia *Fusarium* species (*Fusarium floridanum* and *F. oligoseptatum*) that form nutritional partnerships with *Euwallacea* ambrosia beetles (Aoki et al., 2019, 2018; Kasson et al., 2013). Their strict hydrophobin profiles were identical. SST showed over an order of magnitude more hydrophobins (137) due to a proliferation of group 10 hydrophobins (119 total, both with and without secretion signals). This is an enrichment in this particular hydrophobin group: while approximately 35,000 proteins are predicted in the SST genome, this is not an order of magnitude more proteins than other fungi included in our analysis.

### 5.4 There are filamentous fungi from within Dikarya whose genomes do not appear to encode hydrophobins

Of the 52 proteomes surveyed in the study, 25 including all outgroup organisms failed to recover hydrophobin candidates. Among these are 12 earlier diverging flagellate and non-flagellate fungi, nine members of Dikarya (Basidiomycota and Ascomycota), and four members of Mucoromycota. Within Dikarya, genomes lacking hydrophobin candidates include three obligate biotrophic plant pathogens, including two rust fungi (CQU and PGR, Basidiomycota) and one powdery mildew fungus (ENE; Ascomycota). Five of the remaining six fungal genomes were fungi with a dominant yeast growth form (i.e., *Komagataella pastoris* and *Saccharomyces cerevisiae*), including three human pathogenic species (Basidiomycota: Cryptococcus and Ascomycota: Candida and Pneumocystis). *Athelia termitophila* (ASP), a filamentous fungus that is a termite egg mimic, was the only member of Agaricomycetes included in the study without hydrophobin candidates. The four members of the Mucoromycota lacking canonical hydrophobins included three members of the Mucorales (NSP, RMI, URA), and one Mortierellales (AAM).

### 5.5 Diverse hydrophobin-like proteins (HLPs) are found in fungi and outgroups

Of the sequences contained in the working database, a total of 996 sequences from the 52 proteomes surveyed did not meet the criteria we set for hydrophobin candidacy. Among these sequences, a total of 462 formed 104 groups. Despite their widespread incidence across the complete dataset, it is interesting to note that proteomes with grouped HLPs were restricted to 41 fungi and five outgroups (Figure 8). Four fungal genomes, including two members of Dikarya (PJI and UMA), lacked grouped HLPs. Several of the recovered HLP groups spanned multiple fungal phyla including Group 84, Group 95 and Group 96.

**Figure 8.**
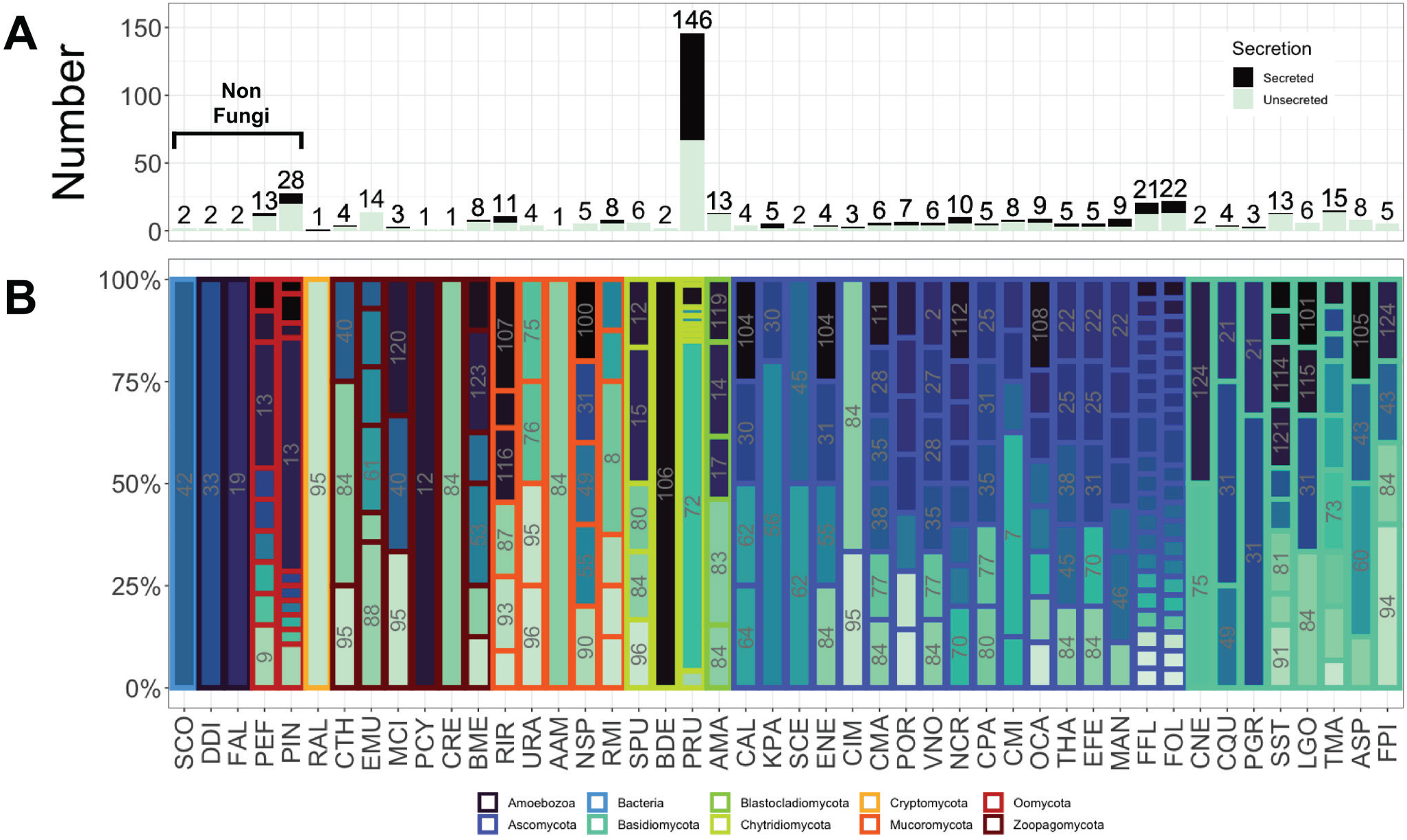
Grouping profiles for hydrophobin-like proteins (HLPs) across organisms, including fungi and non-fungi. (A) Total number of grouped HLPs for each organism is indicated above its respective bar. For each bar the proportion of sequences predicted to contain a secretion signal is indicated by the fill color. (B) Distribution of HLP groups among organisms included in the study. Organisms that did not contain HLP groups and ungrouped HLPs are not included. Labels indicating groups are in grey and are provided for those that account for a minimum of 15% of the total sequences in each respective bar.

Group 84 was present in 20 fungal genomes spanning all included fungal phyla, with the exception of the Cryptomycota. Group 84 represented at least 25% of total HLPs recovered in 5 of 20 genomes. A majority of genomes with Group 84 were in the early diverging fungal lineages and Basidiomycota, with fewer across the Ascomycota. Group 95 was present as a single candidate in ten fungal genomes spanning four phyla, including Cryptomycota, but was noticeably absent across the whole of Basidiomycota. Among these ten genomes, Group 95 represented at least 25% of HLPs recovered from MCI (Zoopagomycota), URA (Mucoromycota), CIM (Ascomycota) and RAL (Cryptomycota). Group 96 was present in five fungal genomes spanning three phyla (Chytridiomycota, Mucoromycota and Basidiomycota).

The remaining groups were found in two or fewer fungal phyla. Of note were Group 100 and Group 120, which were the only groups that included both Oomycota (PEF and PIN) and fungi (NSP in Mucoromycota and MCI in Zoopagomycota, respectively). The other groups from PEF and PIN were exclusively shared, except Group 102 which is unique to PIN. Group 21 and 31 were the only HLPs recovered from both rust genomes (CQU and PGR), both of which lacked hydrophobin candidates. The Hypocreales, which represented about 15% of the sampled genomes in this study, contained 79 grouped HLPs across 28 groups. Two genomes, FOL and FFL, comprised 43 of 79 HLPs from Hypocreales. Surprisingly, Group 84 candidates—one of the most widespread HLP groups recovered in this study—were absent from these two proteomes.

*Pecoramyces ruminantium* (PRU), an anaerobic gut fungus in the *Neocallimastigomycota* that inhabits the rumen and alimentary tracts of multiple ruminant and nonruminant herbivores (Hanafy et al., 2017) contained 146 HLPs spanning 9 distinct groups, none of which were found in any other included proteome (Figure 8). One group, Group 72, accounted for 118 of these HLPs. This observed hyperinflation of HLPs is analogous to that of the hyperinflation of hydrophobin candidates in Group 10 of the artillery fungus *Sphaerobolus stellatus (SST*), considering both the degree of inflation and proliferation of unique other groups within PRU.

### 5.6 Hydrophobin incidence and diversification across fungal lifestyle

The inclusion of diverse fungi spanning various ecologies permitted comparisons of hydrophobin incidence and diversity across a broad array of lifestyles. Fungi that are insect pathogens and insect mutualists were most likely to contain hydrophobin candidates with 4 of 6 (67%) and 3 of 3 (100%), respectively from these groups. A large proportion of plant pathogens and plant mutualists also contained hydrophobin candidates with 5 of 8 (62%) and 3 of 3 (100%), respectively. A lower proportion of saprotrophic fungi and parasitic fungi (including mycoparasites) contained hydrophobin candidates with just 3 of 8 (37.5%) for saprotrophs and 0 of 4 (0%) for parasites. Hydrophobins were completely absent from included proteomes of vertebrate pathogens and mutualists with one exception: the well-known human pathogen and causal agent of Valley Fever, CIM. Known yeasts SCE and KPA reported no hydrophobin candidates.

Regarding diversity, hydrophobin candidates were greatest among saprotrophs (containing 8 different groups; 8 ungrouped candidates), and insect pathogens (6; 10), followed by plant pathogenic fungi (6; 5), insect mutualists (8; 3) and plant mutualists (7; 5). THA contained nearly half of the candidates for plant mutualists (10/23), and SST contributed to over 95% 137/143 of the hydrophobin diversity recovered from saprotrophs (NCR contained 2 and CMA contained 4). The group composition of plant pathogens and insect pathogens mostly overlapped (Groups 92, 69, 79, 117). Groups 5 and 10 were present in plant pathogens and missing from insect pathogens, and Groups 47 and 111 were present in insect pathogens and missing from plant pathogens. Groups 5, 47 and 10 were present in mutualists of both plants and insects. Group 34 was present uniquely in plant mutualists, and Group 26 and 16 were present uniquely in insect mutualists, but not other niches. Among included plant and animal mutualists/pathogens, Group 41 is primarily limited to plant mutualists, but it can also be found among SST candidates.

### 5.7 Principal component analysis (PCA) of the hydrophobin candidate identity matrix reveals a spectrum of sequences and overlapping canonical Class I groups

The variance in the way hydrophobin candidate sequences (and indicator sequences) align and match in a pairwise fashion can be described and visualised by principal component analysis (PCA; Figure 9). More specifically, this is the PCA of a subset of the pairwise sequence identity matrix of the total working sequence database MAFFT. Given these are pairwise comparisons (and so independent), the subset corresponding to the hydrophobin candidates and indicator sequences could be extracted and submitted to PCA. It is important to note, however, that these pairwise comparisons were shaped by the alignment formed from the totality of sequences within the working sequence database. As such, the PCA reflects the structure of the data associated with the pairwise comparison of the hydrophobin candidates and indicator sequences within the context of the broader cysteine-rich working sequence database.

**Figure 9.**
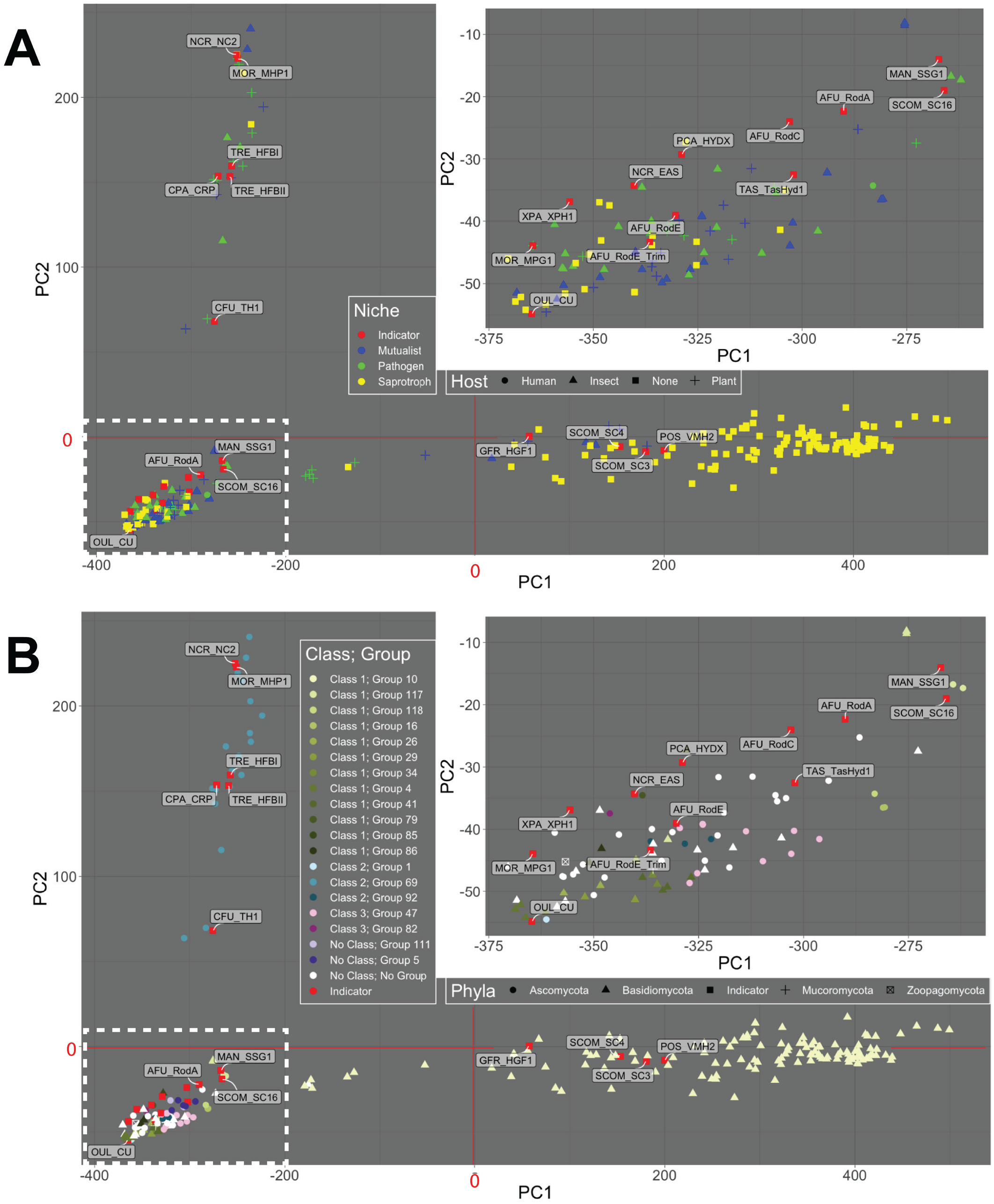
Principal component analysis of hydrophobin candidates from pairwise-identity matrix. PC1 explains 86.9% of variance and PC2 explains 3.45% of the variance. Inset figure in both panels correspond to the region containing the cluster of sequences indicated by the dashed-lined rectangle in the bottom left-hand quadrant of the graph. In panel (A) niche is displayed by marker color and host by marker shape. In (B) the group is indicated by color and the phyla by marker shape. The classes associated with the groups in the figure legend are those developed in this paper (see Section 5.9). In this ontology, Class 2 corresponds to canonical Class II, while Class 1 and Class 3 correspond to canonical Class I. “No Class” refers to sequences that do not belong to any of these classes.

Individual hydrophobin candidate sequences correspond to the data points in **Figure 9** and provide a visualisation of the degree of dispersion of the sequences around the average shared identity represented by the origin of the graph (0,0). The further a sequence lies from the origin, the more significant its differences are from the average. Sequences whose differences from the average are similar cluster together. Interestingly, indicator sequences can be found spanning all regions containing candidate sequences providing confidence for the relevance of the sequences with which they co-cluster. The first two components of the analysis (PC1 and PC2 which are plotted against each other in **Figure 9**) explain 86.9 and 3.45% of the variance in the dataset and separate the data into three distinct regions. It is important to note that the classes presented in **Figure 9** refer to the ontology described in Section 5.9, which we distinguish from the canonical classes by the use of Arabic numerals. Briefly, Canonical Class II corresponds to Class 2 (Arabic numeral) while Canonical Class I corresponds to Class 1 and 3.

Together, the PCA components presented separate the sequences into three distinct regions. More specifically, along the x-axis representing the first component (PC1) the sequences from organisms with a pathogenic niche are limited to the left-hand side (**Figure 9A**) while the right-hand side is limited to Group 10 sequences which are exclusively from Basidiomycotan proteomes (**Figure 9B**). Along the y-axis, in the second component (PC2), Group 69 sequences, which correspond to canonical Class II, separate from the other hydrophobin candidates and are exclusively Ascomycota sequences (**Figure 9A-B**). It is interesting to note that other sequences identified as canonical Class II sequences (i.e., Group 1, Group 92, and the indicator sequence CU) do not cluster with Group 69.

While Group 69 (separated by PC1 and PC2) and Group 10 (separated by PC1) are clearly distinguished in this analysis from other sequences, it is interesting to note that they do not form discrete clusters, but rather display a large range along their respective components. The remaining sequences form a comparatively dense cluster in the bottom left-hand quadrant. The inset figures show this region at a larger magnification revealing overlapping yet distinct groups (**Figure 9B**).

### 5.8 HMM hydrophobin candidate group profiles overlap creating a network that captures the spectrum of hydrophobin sequences

To better understand the spectrum of shared features that contributed to the initial clustering of sequences (**Section 5.7**), we generated an HMM profile representing each hydrophobin candidate group based on their sequences. Given that we purposefully formed these groups with a relatively low threshold (25% sequence identity), we wanted to assess whether group profiles were specific enough to reassign the sequences to their original groupings. Conversely, we were intrigued by the ways in which a particular sequence might fit multiple profiles. As such, we performed a search of all hydrophobin candidates against the library containing the HMM group profiles. The overall workflow for this investigation is illustrated in Figure 5. While most sequences were recognized by multiple profiles, assignment to the top scoring sequence model resulted in group assignments that were almost identical to initial assignments (98%; 203 of the 207 candidate hydrophobins with groups). The four sequences with inconsistent groups were assigned to multiple similarly ranked profiles suggesting these few instances reflect overlaps among group profiles.

Of the 31 hydrophobin candidates that were not captured in the initial groupings, 21 were assigned existing groups by the HMM profiles. Noteworthy sequences among these include a SST protein (SST_KIJ24675.1; a class II candidate in Basidiomycota) classified into Group 1 (containing OUL_CU) and one protein each from CPA (CPA_KAF3767202.1) and CMI (CMI_XP_006672063.1) classified into group 118 (previously occupied by only CIM). A reclassified UMA protein (UMA_XP_011391376.1) was placed into Group 117, which provides further support for this cross-phyla group. Additionally, a POR protein (POR_XP_003716609.1) was placed into Group 34, suggesting this group also has cross-phyla features. It is important to note that these adjustments do not significantly impact the results or interpretation of the original groupings previously described.

Most group HMM models contained a core 8-cysteine motif. These were specifically all Class II groups, and Groups 10, 117, 34, 85, 79, 47, 118 and 82. Groups 4 (4 cysteines; the fewest), 29 (7), 5 (7), 26 (6), 86 (6), 111 (6) and 41 (5) had patterns with fewer cysteines, and the Groups 16 model contains a pattern consisting of 9 cysteines.

When considering the overall models, however, each hydrophobin candidate group HMM profile varies in its breadth and the ways in which they overlap with each other. While each profile highlights features that unite sequences in a given group, each profile also accommodates varying degrees of sequence diversity. To better understand these features, we established pairwise co-classification: the ability of two groups to overlap in terms of the sequences which they contain. We quantified and visualised co-classification of hydrophobin candidate sequences into their hydrophobin candidate Group and canonical Classes (Pfam assigned) in the form of a pairwise co-classification heat map (**Figure 10**). The extent to which sequences co-classify between groups varies widely. Within the matrix lies two clusters of groups with a high degree of co-classification including 100%. At the core of two of these clusters are the sequences which are assigned canonical class I and II Pfam profiles. More specifically, looking along the X axis of **Figure 10**, all sequences contained in Groups 4 through 79 as well as group 118 co-classify entirely with the canonical Class I. The sequence content of Groups 47 and 5, however, only partially overlap with canonical Class I. Conversely, not all sequences contained in groups defined as belonging to canonical Class I were identified by the canonical Pfam Class I model. This was to be expected given the method chosen to associate canonical Classes to the candidate Groups developed in the study (as described in Section 5.2). In contrast, all Class II hydrophobin co-classify highly with one another (i.e., ranging from 92-100% overlap) and do not co-classify (i.e., less than 3% across only 3 groups) with any groups associated from canonical Class I. As such, the Class II cluster (Groups 69, 1, 92 and Pfam Class II) can be seen as resolved from the other groups.

**Figure 10.**
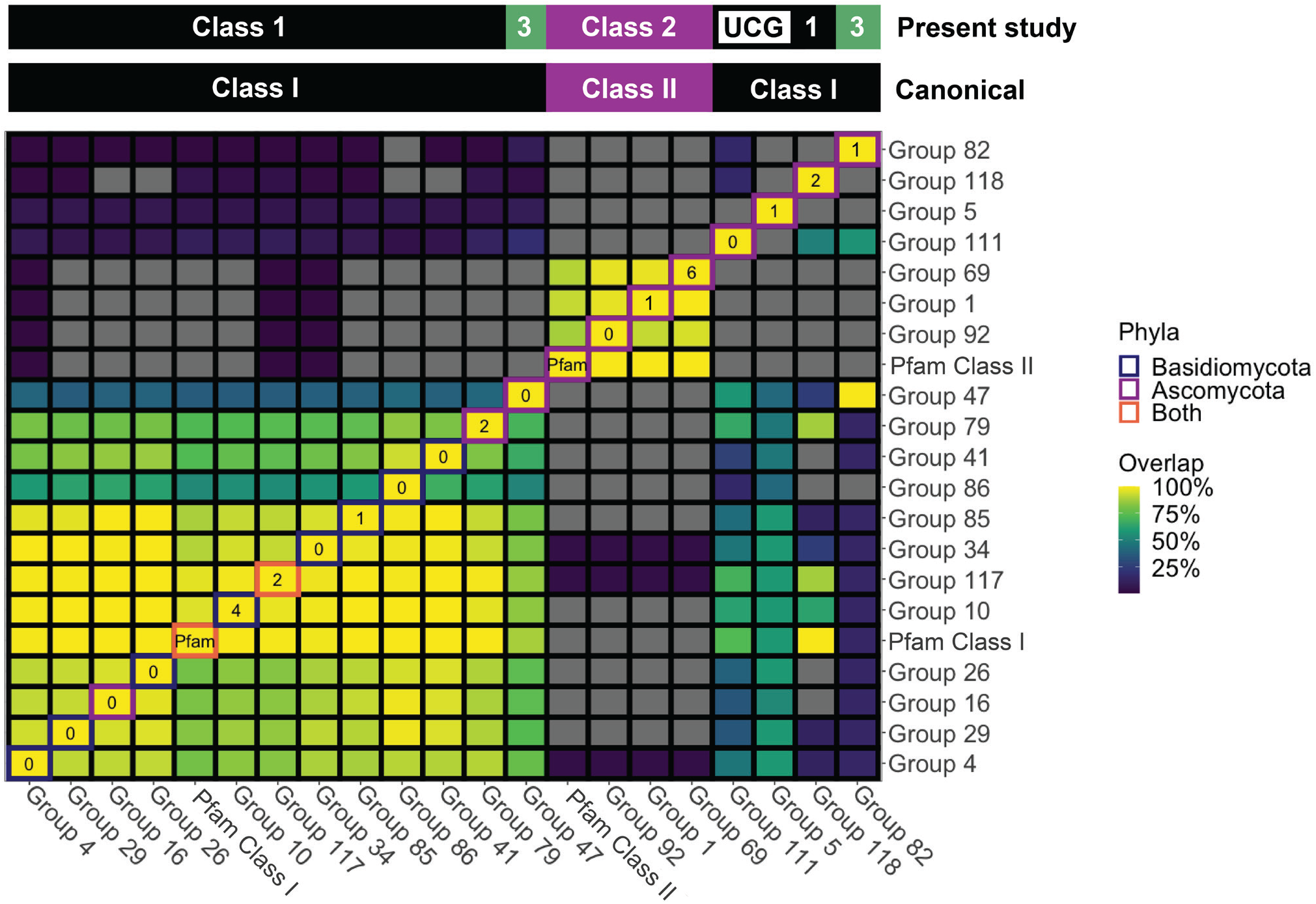
The pair-wise comparison of sequences recognized by each model (co-classification) reveals varying degrees of overlap for hydrophobin candidate Group sequence models established in the study as well as canonical Pfam models. The extent of overlap between the sequences recognized by two given models are reported according to the column (i.e., percent of members recognized by columnar group that were also recognized by the row-wise group). Gray boxes signify no overlap in group content. Boxes along the diagonal of the matrix are used to annotate the number of indicator sequences in each columnar group.

While this co-classification clearly recapitulates the canonical Class I and Class II dichotomy, it also highlights ways in which the candidate groups may be connected across the natural spectrum of hydrophobin sequences. For instance, class I sequences co-cluster with class II sequences in groups 34, 117, and 4 (less than 3%). Two hydrophobin candidates, both from Trichoderma co-classified into these groups. Additionally, one candidate from Trichoderma (THA_XP_024780278-1) co-classifies into Class I Group 34 and 4, as well as all Class II groups, including the canonical Class II category. Intriguingly, the Class II indicator sequence TRE_HFBII co-classified into all Class II groups, but also the Class I Group 117, which is a cross-phyla Class I-containing group.

Within Group 4 lies another interesting potential connection between Class I and Class II sequences. Group 4 consists of four SST proteins.These sequences align in a staggered fashion resulting in an HMM profile that only captures 4 of the cysteines in the 8-Cys pattern present in each. Of these, SST_KIJ24675.1 co-classifies with the Class II Group 1 despite Class II sequences having been canonically defined as found exclusively in Ascomycota. This is the only instance in our study that suggests a basidiomycotan sequence might be a Class II sequence. Furthermore, this sequence syntenously spans Group 4 candidates. As such, this unique SST group may represent a transition between classes: the truncated Group 4 model appears to be sensitive to similarities between groups.

### 5.9 The co-classification of sequences provides an ontological framework that can be used to contextualise the spectrum of hydrophobin candidate sequences captured by the survey

The co-classification matrix suggests three overlapping regions of hydrophobin candidate sequence space (i.e., all combinations of amino acids that we have defined as hydrophobin candidates in this study). These regions were determined bioinformatically by creating by binning of groups which share 100% overlap with another group into classes: this was done by adapting our sequence similarity network approach to calculated overlaps between groups. To be congruent with the canonical subdivision of sequences into Class I and Class II, we have numbered these regions with Arabic numerals as Class 1, Class 2, and Class 3. These are summarized in Figure 11 along with their relationships with the canonical classes and hydrophobin candidate groups developed in this study. Largely, canonical Class II as defined by Pfam corresponds to Class 2, while Class I defined by Pfam corresponds to Class 1 and 3. It is important to note that within Class 3, Group 82, which consists of indicator sequence EAS and its homolog are not recognized by the canonical hydrophobin Pfam model.

**Figure 11.**
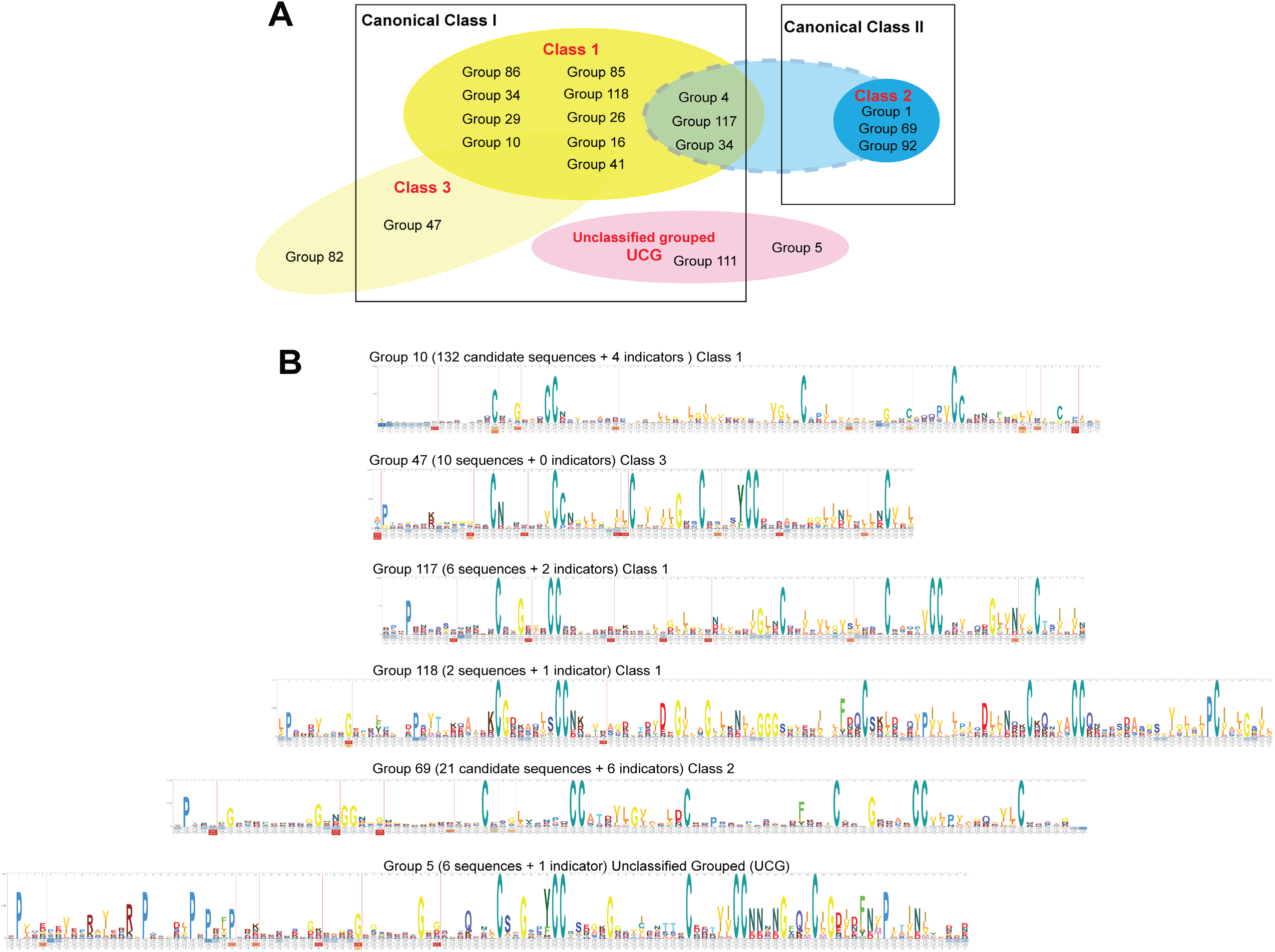
HMM established for each group accommodate a range of degrees of variability. Some models are more specific than others. As such, groups can be seen as overlapping in multiple ways (A). Through this lens the top-down relational hierarchy frequently found in phylogenetic studies is not imposed. (B) Examples of models for groups described in (A).

Describing the survey within the context of this ontology, many more Class 1 hydrophobin candidate sequences were uncovered compared to the Ascomycota-specific Class 2 hydrophobin candidates, despite the inclusion of twice as many Ascomycota proteomes (18) compared to the next most abundantly sampled fungal phylum, Basidiomycota (9). *Sphaerobolus stellatus* (Basidiomycota), accounted for 129 of the 163 Class 1 hydrophobins reported. Hydrophobins from just three additional Basidiomycota (*Fomitopsis pinicola, Leucoagaricus gongylophorus*, and *Tricholoma matsutake*) accounted for >75% of the remaining Class 1 hydrophobins. Class 1 candidates from Ascomycota were single candidates from CIM, CMI, FFL, FOL and POR and two candidates from MAN. Class 3 hydrophobins (Group 47 and Group 82) consisted of 11 candidates from 8 Ascomycota genomes. Group 47 represented a majority of the Class 3 hydrophobins and were predominantly from hypocrealean genomes, with the exception of *Corollospora maritima* (Microascales). Group 82 contained a single hydrophobin from Neurospora crassa, which matched an indicator hydrophobin (EAS) from this same species, which has been intensively studied.

Unclassified grouped (UCG) hydrophobins consisted of 8 hydrophobins comprising two groups (Group 5 and Group 111) from 6 Ascomycota genomes that did not display 100% overlap with any other group. Both of these groups had no overlap with Class 2 groups and low overlap with various Class 1 groups. Group 5 represented a majority of these UCG hydrophobins and included the indicator TasHyd1, a hydrophobin from the biocontrol agent *Trichoderma asperellum* that is involved in plant root colonization (Viterbo and Chet, 2006). The genomes containing UCG and Class 3 proteins noticeably overlapped (5 of 9 total proteomes containing these groups; OCA, FFL, FOL, THA, and CMA). POR candidates were present in UCG and not Group 3, and MAN, EFE and NCR contained proteins in Group 3 and not UCG. Unclassified Ungrouped (UCU) hydrophobins comprised the remaining 31 strict hydrophobins recovered. UCU hydrophobins spanned four fungal phyla (Ascomycota, Basidiomycota, Mucoromycota and Zoopagomycota). UCU candidates were primarily found in Ascomycota (15 candidates) and Basidiomycota (14), with one candidate each from Mucoromycota and Zoopagomycota.

## 6. Discussion

Our proteomic survey, supported by both established and recent literature, expands the known repertoire of hydrophobins and contributes to the emerging view of this protein family. In contrast to previous efforts, we chose to center fungal ecology and put aside questions of evolutionary origin to establish a richly annotated library of hydrophobin sequences. This allowed us to re-evaluate the canonical definition of the hydrophobin family and its Class I and II subdivisions and build an ontology, based on our modern understanding of the family, that captures the remarkable sequence diversity and structural plasticity of this protein family within their ecological context.

### Members of the hydrophobin protein family are not, a priori, hydrophobic

First, it is of particular importance to address what can be seen as the misleading nomenclature of the term “hydrophobin” in light of our modern understanding of the mechanisms underlying their assembly and resulting functions. The term “hydrophobin” was first coined by (Rosenberg and Kjelleberg, 1986) as a category for biomolecules thought to associate with the exterior of the bacterial cell and contribute to its decreased wettability (referred to in their work as hydrophobicity) without ascribing a specific biological function to the phenomenon. This hydrophobicity, however, is not a property of the individual biomolecules; rather, it is an emergent property of the association of the hydrophobin component with the cellular surface and surrounding microenvironment. In other words, in nature, the ability of a surface to repel water is most often the result of a number of contributing factors both chemical (i.e. the tendency of nonpolar amino acids in the protein to be excluded by water – a polar solvent); and physical (i.e. nanoscopic and/or microscopic structures that decrease the ability of a droplet of water to adhere to a surface) (Yoshimitsu et al., 2002). Through this lens, proteins belonging to the hydrophobin family need not be overly hydrophobic at the molecular scale prior to assembly in order to contribute to the hydrophobicity of a particular surface or interface. Rather, the hydrophobicity that is discussed in such a context emerges from the assembly of biomolecular components within the dynamic context of the ecosystem and can involve significant rearrangements of molecular structures. This perspective is of particular importance to both capturing and understanding the rich sequence diversity found in the hydrophobin protein family. Furthermore, our review of the literature clearly establishes that hydrophobins modify the wettability of surfaces, not only increasing wettability, but decreasing it too (Cai et al., 2020).

### Hydrophobin function is driven by assembly in the ecosystem that relies, to varying degrees, on specific molecular interactions

As summarized in (Table 1), it is well established that hydrophobins do not share a single unifying biological function. Rather, they are most frequently described as providing a palette of properties (Zampieri et al., 2010) for the modification of surfaces and interfaces (Linder, 2009), and play multiple roles in fungal biology (Wösten, 2001). We propose the following complementary view: hydrophobin function can be described as driven by assembly within an ecosystem to fulfill a material function (e.g., coating, adhesion, sensing). In contrast to many functions that rely on highly specific amino acid interactions at the molecular scale, a wide range of material functions are a result of more general physicochemical properties. For instance, most proteins with an amphiphilic nature (e.g., charge segregation or hydrophobic patch at their surface) can accumulate at fluid interfaces (e.g., air-liquid interface) and decrease the surface tension. This general property does not rely on the conservation of specific amino acid interactions, but rather the overall amphiphilic chemical character of the biomolecule. As such, many highly expressed hydrophobin proteins can modulate surface tensions when secreted into the surrounding environment. On the other hand, fungi have also evolved unique hydrophobins whose properties, once assembled, critically rely on specific interactions. For instance, the remarkable elasticity of the interfacial films formed by the Class II hydrophobin HFBI (T. reesei; study indicator)(Cox et al., 2007), relies on highly specific intermolecular amino acid interactions absent in other hydrophobins. These specific interactions lead to the precise geometric packing of the protein (Szilvay et al., 2007) and resulting chemical interactions (Lienemann et al., 2013)(Szilvay et al., 2007) critical for thermodynamics that highly favor this unique assembled state (Paananen et al., 2021).

### The hydrophobin family is best defined as a generic molecular structure capable of accommodating remarkable structural and chemical diversity. The family’s underlying amino acid sequence diversity can be captured in a spectrum-based ontology (co-classification)

As we describe in Section 3 (Figure 2), an emerging definition of the hydrophobin family is based on the generic features of its shared molecular structure that accommodates significant regions of chemical and structural diversity. The plasticity accommodated by the protein’s core fold both unifies the family and is the basis for the challenges faced by existing alignment-based hydrophobin ontologies. For instance, alignment-based approaches are of limited use for understanding the ways in which these proteins are phylogenetically related to each other across the fungal kingdom given the absence of long spans of sequence similarity (Mgbeahuruike et al., 2013) (Littlejohn et al., 2012)(Plett et al., 2012). There is significant value in describing hydrophobin sequence-space to better draw connections between the amino acid sequence variability and the wide spectrum of solution properties and molecular assembly mechanisms captured by this family. We have previously proposed (Gandier, 2017)(Gandier and Master, 2018)(Gandier et al., 2017) that to better understand both the relationship between sequence and biological function, the family ought to be understood through the lens of a spectrum rather than discrete categories. Herein we further hypothesized that this observed spectrum has been shaped by fungal ecology linking hydrophobin amino acid sequence and the fungal environment. Hydrophobins exemplify that fungi are not passive recipients of their surroundings. To appreciate the spectrum of hydrophobin sequence diversity and pleiotropic functions, the dynamic fungal environment—determined and driven by fungal ecology—must be considered.

To achieve this, a critical literature review was conducted to identify the slate of indicator sequences that would help guide our analysis by providing us with the bounds of what is known about the experimentally validated properties of this protein family. These indicators can inform hypotheses on function, regulation and/or structure of group members to varying degrees based on sequence identity. Overall, the relationships among indicators in our analysis recapitulated known class delineations, providing additional support for our approach. Our pairwise co-classification placed these indicators in context of the observed spectrum of hydrophobin sequences. From across the spectrum of hydrophobins within our analysis, we identified three discernable regions (or classes) of hydrophobins, in addition to hydrophobins that could not be further classified. Our identified classes (Class 1, 2 and 3) largely recapitulate canonical Class I and Class II delineations, but provide more resolution and uncover undescribed relationships among hydrophobins. Class 3 is composed solely of Group 47 and Group 82 (containing EAS). Three groups of Class 1 hydrophobins included candidates from both Basidiomycota and Ascomycota: Group 117 boasts indicators from both phyla. Only 8 proteins across Groups 5 (FFL, FOL, THA, POR and CMA) and 111 (limited to OCA) were unclassified by our approach: these groups may have evolved novel functions among these ascomycotan fungi.

### Hydrophobins are not limited to Dikarya

Overall, the survey was designed to examine the phylogenetic breadth of hydrophobins with probing inclusion of multiple relevant outgroups and early divergent fungi, which have not yet been included in such studies. Overall, it spanned eight fungal phyla and more than 25 orders, uncovering 238 hydrophobin candidates from across 20 fungal proteomes. While broadly reported as limited to filamentous fungi in Dikarya, members of the hydrophobin protein family have been identified in the recently sequenced genomes of *Caulochytrium protostelioides* (Chytridiomycota) and Mortierella elongata (Mortierellamycotina: Mucoromycota) (Ahrendt et al., 2018). Our own survey has identified an additional hydrophobin in Mucoromycota, specifically the Glomeromycotina (*Rhizophagus irregularis*), and expands the hydrophobin family into the phylum Zoopagomycota (*Entomophthora muscae*). This supports recent speculation that these proteins are involved in spore adhesion in E. muscae (Elya and De Fine Licht, 2021). Plainly, our survey supports the emerging view that hydrophobins are not limited to Dikarya. An additional 996 hydrophobin-like proteins (HLPs) were identified across all fungal proteomes, with the exception of Microsporidia. HLPs were additionally found from Oomycota, Amoebozoa, and bacteria. It is important to note that all HLPs identified were done so through the permissive screen for the 6-Cys pattern. As such, they are anticipated to contain a wide range of sequences including those that are functionally and structurally unrelated to hydrophobins. While outside the scope of this study, investigating how hydrophobin sequences relate to such other small, secreted cysteine-rich proteins could help fill the gaps in our understanding of how various cysteine patterns emerged over the course of fungal evolution.

### Hydrophobins may play intracellular functions

Our analysis uncovered few, but widespread hydrophobin candidates whose sequences were not predicted to contain a secretion signal. Interestingly, these do not form separate sequence groupings; rather, these group together with hydrophobin candidates containing a predicted secretion signal. The proliferation of Group 10 hydrophobins in SST exemplifies this apparent relationship between the presence and lack of secretion signal. Among this group, 48.7% contain predicted secretion signals, while the remaining 51.3% do not. Although the biological functions of hydrophobins have been considered exclusively as extracellular, a recent study (Cai et al., 2021) described the formation of tonoplast-like structures in *Trichoderma* species, enriched with hydrophobins, that align within hyphae before secretion. These authors propose that prior to secretion, hydrophobins may form structures that play roles in maintaining turgor pressure in aerial hyphae to support, for instance, the formation of spores. While such roles must be further investigated, this observation underlines the pleiotropic nature of hydrophobins and provides a clear example in which these proteins impact fungal fitness and cellular structure in ways that have yet to be fully explored. The discovery of widespread hydrophobins herein that were not predicted to contain secretion signals suggests that hydrophobins may play an unappreciated intracellular role that ought to be further investigated. In light of the (Cai et al., 2021) study, it could be speculated that, for instance, certain mechanical properties of fungi that rely on turgor pressure, such as appressoria formation (Ryder et al., 2022), could be tuned via intracellular accumulation of hydrophobins. As such, the potential intracellular roles hydrophobins play ought to be further studied in the context of fungal development.

### Ecological drivers of hydrophobin diversity

The convincing role hydrophobins play for aerial hyphae and spore production has led to the view that hydrophobins are limited to filamentous fungi—or that hydrophobins are not present in yeasts. The categories yeasts and filamentous fungi are distinguished by distinct development and ecology. Five yeasts included in this study (*Candida albicans*, *Cryptococcus neoformans*, *Komagataella pastoris*, *Pneumocystis jirovecii*, and *Saccharomyces cerevisiae*) lacked hydrophobin candidates, which likely reflects their specialized ecology of growth via budding, not shared evolutionary history (Nagy et al., 2014). Yeasts are paraphyletic, and we included yeasts spanning three classes across two fungal phyla. Interestingly, multiple filamentous fungi included in this study also lack hydrophobin candidates (Section 5.4). These fungi were notably engaged in specialized symbioses as rusts, powdery mildew or, in one fungus that acts as a termite egg-mimicking symbiont: the only filamentous basidiomycete without any hydrophobin candidates. Depending on ecological need, specialized fungal lifestyles are marked by shifted hydrophobin content: either through loss or diversification of these proteins for new contexts. For example, their diversification and duplication in some ecological contexts, such as in the saprotrophic and necrotrophic phytopathogenic fungi is notable.

Distribution of Groups followed distinct patterns based on ecology. Groups 5 (among Ascomycota) and 10 (Basidiomycota) were found in plant pathogens. These groups were not found in insect pathogens, which instead contained Groups 47 or 111. Groups 5 and 47 (among Ascomycota) and 10 (among Basidiomycota) were present in mutualists of plants and insects, with additional strict groups limited to insect mutualists (e.g., Group 16 in *Fusarium* species) or plant mutualists (e.g., Group 34 in *Tricholoma matsutake*). Group 5 is associated with plant mutualism in Sordariomycetes, including a marine mutualist CMA(Cuomo et al., 1985). This marine plant mutualist presents an opportunity to study self-assembly dynamics at the host-mutualist interface under high salt conditions. Group 5 notably contains TasHyd1, which has been implicated in root colonization (Viterbo and Chet, 2006).

*Trichoderma* seems to have unique features among organisms included in this study that may bridge the varied ecological niches of *Trichoderma* ranging from root mutualist to mycoparasite. The *Trichoderma harzianum* genome in particular has an unusually large and diverse repertoire of hydrophobins (11), rivaled only by SST (137) in which Group 10 sequences are amplified, and LGO (17). Interestingly, one of Trichoderma’s hydrophobin candidate groups (Group 1) with cerato ulmin (CU from *Ophiostoma*). The expansive repertoire of hydrophobins in *Trichoderma* does not reflect the reality in most other fungi.

This diverse set of hydrophobins may present opportunities to understand fungi that engage in different lifestyles, such as *Ophiostoma*. Another vascular wilt pathogen included in this study, VNO, lacked Group 1 hydrophobins, but did share a somewhat rare Group 92 hydrophobin with *Trichoderma*. *Trichoderma* hydrophobin candidates were found to span classes, specifically Groups 34 and 4.

*Coccidioides immitis* (CIM) is a human pathogen known to cause Valley Fever in the Southwestern United States. This fungus is soil-borne and causes infection through inhalation. We were surprised to find that this pathogen contained a strict hydrophobin that grouped exclusively with our indicators RodA and RodC (Group 118). RodA is an A. fumigatus virulence factor that masks Dectin-1 and Dectin-2 recognition of conidia (Aimanianda et al., 2009)(Carrion et al., 2013). As soil-borne fungi causing lung infections, both of these ascomycete human pathogens are linked by their ecology. Our application of the Group 118 HMM profile on all candidates revealed that additional CPA and CMI candidates are recognized by this model: both of these fungi are pathogens (plants and insects, respectively).

### Specific hydrophobin groups underlie major divisions of fungi

Group patterns additionally followed known phylogenetic relationships. Group 10 was present in nearly all included Basidiomycota, and Group 69 was present in nearly all included Ascomycota. Members from these clades without these groups have specialized lifestyles accompanying their altered hydrophobin repertoire. While Group 10 is present across basidiomycota, SST contains an order of magnitude more Group 10 candidates. This is apparently due to a proliferation of these genes within the genome of the cannonball fungus, named for its ability to eject a bolus from woody substrate that sticks on contact to deliver spores. The two most closely related fungi in our analysis were both *Fusarium* species, and their hydrophobin profiles resolved highly with identical strict hydrophobin profiles and near identical HLP profiles. This suggests that our method can resolve a high amount of diversity among cysteine-rich proteins when applied to fungi from the same genus. The hydrophobin candidates from early divergent fungi (EMU and RIR) were both ungrouped, which is not too surprising considering studies on hydrophobins have largely been limited to Dikarya. In contrast, most Sordariomycetes contained multiple ungrouped strict hydrophobin candidates, including well-studied fungi like THA. These ungrouped candidates might represent evolution of hydrophobins that are unique to a particular niche or clade, but their function and phylogenetic limits must be tested by further studies.

### Hydrophobin-like proteins (HLPs) share a distinct cysteine pattern within their primary amino acid structure and their presence varies across fungi

As anticipated, HLPs are certainly composed of multiple protein families serving diverse purposes. They are, however, unified by the presence of our 6-Cys pattern screening criteria and therefore represent a particular subset of cysteine-rich proteins encoded within the genomes of the organisms included in our study. The groups formed by the HLPs also show patterns that can be linked variously to ecology, with certain groups strongly conserved and spanning all or most sampled fungal phyla. There were also a number of clear patterns in their distribution. For instance, Group 100 and 120 spanned early divergent fungi and Oomycota, which provides a link between phylogenetically distinct groups that share underlying development and ecology. While the rust fungi CQR and PGR lacked hydrophobin candidates, both contained HLP groups 21 and 31, which themselves were limited to these rusts. Two HLPs were identified from CIM (Group 84 and Group 95), which were widespread across Kingdom Fungi. These other fungi may provide a tractable model for understanding the role of Group 84 and Group 95 candidates during CIM development. Among hydrophobin candidates, SST stood out with an extreme proliferation within Group 10, and PRU revealed a similar pattern within HLP Group 72. Unlike Group 10, which was a widespread hydrophobin candidate group among Basidiomycota, this hyper-inflated PRU group was not found from other fungi, including other chytrids. Though their environments and developmental stages are quite different, SST and PRU both are subjected to a range of host substrates in mulch or ruminant guts, respectively. Direct investigation into these groups among PRU and SST is required, but we hypothesize that these instances of hyper-inflation may serve as a cradle for innovation by providing variety where neofunctionalization can occur before the observed massive redundancy is lost. A similar process may have led to the many ungrouped or unique groups observed in our analysis.

## 6. Conclusions and perspectives

Altogether, our survey of hydrophobin protein diversity across Kingdom Fungi has revealed that ecology is a major factor driving the family’s remarkable spectrum of amino acid sequence chemistry. We found that fungi have distinct profiles of hydrophobin classes and groups that share, to varying extents, sequence features whose contribution to molecular and biological function can now be experimentally explored within the framework developed. While our ontology encompasses the canonical Class I and Class II family subdivisions, it provides a novel feature: the co-classification of sequences as a means of describing hydrophobins as a spectrum of overlapping features (i.e., no enforcement of discrete categories) without the imposed top-down relational hierarchy often found in phylogenetic studies. Such an approach will be critical to describing and understanding the sequence variability that could drive function within the hydrophobin family. For instance, it could be speculated that fungi (and many other organisms) are able to sense and behaviorally interpret hydrophobin profiles of other fungi—an ecological role for this protein family that could be driven by the variability of their protein chemistries. Such interactions could occur directly via associations with hydrophobins or indirectly via changes in the physicochemical properties of the surrounding environment driven by hydrophobin assembly, as suggested by the in vitro study of MPG1 and MHP1 of Magnaporthe grisea (Pham et al., 2016). As such, our proposed ontology could provide a framework to design the studies that will explore the molecular grammar contained within the variable regions of hydrophobin sequences that may drive what can be seen as material assembly.

The process of hydrophobin material assembly is inextricable from fungal ecology: from filamentous growth to the establishment of fundamental symbioses. In other words, hydrophobins do not act in isolation. In this context, our work provides a modern view of the hydrophobin family and highlights their complexity. To unlock the unexamined potential of hydrophobins will require continued interdisciplinary collaboration.

## Supporting information

Supplemental file containing 3 tables.

## 7. Acknowledgements

This work was a part of the HFBMat project funded by the Novo Nordisk Foundation Postdoctoral Fellowship for “Research within Biotechnology-Based Synthesis & Production”. We are grateful to Dr. Joey Spatafora for providing permission to include Corollospora maritima CBS 119819 (CMA) sequence data in our analysis. These data were produced by the US Department of Energy Joint Genome Institute (https://ror.org/04xm1d337; operated under Contract No. DE-AC02-05CH11231) in collaboration with the user community. Mention of trade names or commercial products in this publication is solely for the purpose of providing specific information and does not imply recommendation or endorsement by the United States Department of Agriculture. USDA is an equal opportunity provider and employer.

## References

Ahrendt SR, Quandt CA, Ciobanu D, Clum A, Salamov A, Andreopoulos B, Cheng J-F, Woyke T, Pelin A, Henrissat B, Reynolds NK, Benny GL, Smith ME, James TY, Grigoriev IV. 2018. Leveraging single-cell genomics to expand the fungal tree of life. Nat Microbiol 3:1417–1428.

Aimanianda V, Bayry J, Bozza S, Kniemeyer O, Perruccio K, Elluru SR, Clavaud C, Paris S, Brakhage AA, Kaveri SV, Romani L, Latgé J-P. 2009. Surface hydrophobin prevents immune recognition of airborne fungal spores. Nature 460:1117–1121.

Almagro Armenteros JJ, Tsirigos KD, Sønderby CK, Petersen TN, Winther O, Brunak S, von Heijne G, Nielsen H. 2019. SignalP 5.0 improves signal peptide predictions using deep neural networks. Nat Biotechnol 37:420–423.

Aoki T, Kasson MT, Berger MC, Freeman S, Geiser DM, O’Donnell K. 2018. Fusarium oligoseptatum sp. nov., a mycosymbiont of the ambrosia beetle Euwallacea validus in the Eastern U.S. and typification of F. ambrosium. Fungal Systematics and Evolution 1:23–39.

Aoki T, Smith JA, Kasson MT, Freeman S, Geiser DM, Geering ADW, O’Donnell K. 2019. Three novel Ambrosia Fusarium Clade species producing clavate macroconidia known (F. floridanum and F. obliquiseptatum) or predicted (F. tuaranense) to be farmed by Euwallacea spp. (Coleoptera: Scolytinae) on woody hosts. null 111:919–935.

Beever RE, Dempsey GP. 1978. Function of rodlets on the surface of fungal spores. Nature 272:608–610.

Bell-Pedersen D, Dunlap JC, Loros JJ. 1992. The Neurospora circadian clock-controlled gene, ccg-2, is allelic to eas and encodes a fungal hydrophobin required for formation of tlie conidiaI rodlet layer. Genes & Development 6:2382–2394.

Bentley SD, Chater KF, Harper D, Bateman A, Brown S, Chandra G, Chen CW, Collins M, Cronin A, Fraser A, Goble A, Hidalgo J, Hornsby T, Howarth S, Rutter S, Seeger K, Saunders D, Sharp S, Squares R, Squares S, Taylor K, Warren T, Wietzorrek A, Woodward J, Barrell BG, Parkhill J, Hopwood DA. 2002. Complete genome sequence of the model actinomycete Streptomyces coelicolor A3(2). Nature 417:141–147.

Black SD, Mould DR. 1991. Development of hydrophobicity parameters to analyze proteins which bear post- or cotranslational modifications. Analytical Biochemistry 193:72–82.

Cai F, Gao R, Zhao Z, Ding M, Jiang S, Yagtu C, Zhu H, Zhang J, Ebner T, Mayrhofer-Reinhartshuber M, Kainz P, Chenthamara K, Akcapinar GB, Shen Q, Druzhinina IS. 2020. Evolutionary compromises in fungal fitness: hydrophobins can hinder the adverse dispersal of conidiospores and challenge their survival. ISME J 14:2610–2624.

Cai F, Zhao Z, Gao R, Chen P, Ding M, Jiang S, Fu Z, Xu P, Chenthamara K, Shen Q, Bayram Akcapinar G, Druzhinina IS. 2021. The pleiotropic functions of intracellular hydrophobins in aerial hyphae and fungal spores. PLoS Genet 17:e1009924.

Carpenter CE, Mueller RJ, Kazmierczak P, Villalon DK, Van Alfen NK. 1992. Effect of a virus on the accumulation of a tissue-specific cell-surface protein of the fungus cryphonectria (endothia) parasitica. Molecular Plant-Microbe Interactions 4:55–61.

Carrion S de J, Leal SM, Ghannoum MA, Aimanianda V, Latgé J-P, Pearlman E. 2013. The RodA Hydrophobin on Aspergillus fumigatus Spores Masks Dectin-1- and Dectin-2–Dependent Responses and Enhances Fungal Survival In Vivo. JI 191:2581–2588.

Chang Y, Wang S, Sekimoto S, Aerts AL, Choi C, Clum A, LaButti KM, Lindquist EA, Yee Ngan C, Ohm RA, Salamov AA, Grigoriev IV, Spatafora JW, Berbee ML. 2015. Phylogenomic Analyses Indicate that Early Fungi vEvolved Digesting Cell Walls of Algal Ancestors of Land Plants. Genome Biology and Evolution 7:1590–1601.

Chang Y, Wang Y, Mondo SJ, Ahrendt S, Andreopoulos W, Barry K, Beard J, Benny G, Blankenship S, Bonito G, Cuomo CA, Desirò A, Gervers KA, Hundley H, Kuo A, LaButti K, Lang BF, Lipzen A, O’Donnell K, Pangilinan J, Reynolds N, Sandor L, Smith MW, Tsang A, Grigoriev IV, Stajich J, Spatafora JW. 2022. Fungi Are What They Secrete: Evolution of Zygomycete Secretomes and the Origins of Terrestrial Fungal Ecologies. SSRN Journal.

Copp JN, Akiva E, Babbitt PC, Tokuriki N. 2018. Revealing Unexplored Sequence-Function Space Using Sequence Similarity Networks. Biochemistry 57:4651–4662.

Cox AR, Cagnol F, Russell AB, Izzard MJ. 2007. Surface Properties of Class II Hydrophobins from Trichoderma reesei and Influence on Bubble Stability. Langmuir 23:7995–8002.

Crouch JA, Dawe A, Aerts A, Barry K, Churchill ACL, Grimwood J, Hillman BI, Milgroom MG, Pangilinan J, Smith M, Salamov A, Schmutz J, Yadav JS, Grigoriev IV, Nuss DL. 2020. Genome Sequence of the Chestnut Blight Fungus Cryphonectria parasitica EP155: A Fundamental Resource for an Archetypical Invasive Plant Pathogen. Phytopathology® 110:1180–1188.

Cuomo V, Vanzanella F, Fresi E, Cinelli F, Mazzella L. 1985. Fungal flora of Posidonia oceanica and its ecological significance. Transactions of the British Mycological Society 84:35–40.

De Schutter K, Lin Y-C, Tiels P, Van Hecke A, Glinka S, Weber-Lehmann J, Rouzé P, Van de Peer Y, Callewaert N. 2009. Genome sequence of the recombinant protein production host Pichia pastoris. Nat Biotechnol 27:561–566.

De Vries OMH, Moore S, Arntz C, Wessels JGH, Tudzynski P. 1999. Identification and characterization of a tri-partite hydrophobin from Claviceps fusiformis – A novel type of class II hydrophobin. Eur J Biochem 262:377–385.

Dean RA, Talbot NJ, Ebbole DJ, Farman ML, Mitchell TK, Orbach MJ, Thon M, Kulkarni R, Xu J-R, Pan H, Read ND, Lee Y-H, Carbone I, Brown D, Oh YY, Donofrio N, Jeong JS, Soanes DM, Djonovic S, Kolomiets E, Rehmeyer C, Li W, Harding M, Kim S, Lebrun M-H, Bohnert H, Coughlan S, Butler J, Calvo S, Ma L-J, Nicol R, Purcell S, Nusbaum C, Galagan JE, Birren BW. 2005. The genome sequence of the rice blast fungus Magnaporthe grisea. Nature 434:980–986.

Druzhinina IS, Chenthamara K, Zhang J, Atanasova L, Yang D, Miao Y, Rahimi MJ, Grujic M, Cai F, Pourmehdi S, Salim KA, Pretzer C, Kopchinskiy AG, Henrissat B, Kuo A, Hundley H, Wang M, Aerts A, Salamov A, Lipzen A, LaButti K, Barry K, Grigoriev IV, Shen Q, Kubicek CP. 2018. Massive lateral transfer of genes encoding plant cell wall-degrading enzymes to the mycoparasitic fungus Trichoderma from its plant-associated hosts. PLoS Genet 14:e1007322.

Eichinger L, Pachebat JA, Glöckner G, Rajandream M-A, Sucgang R, Berriman M, Song J, Olsen R, Szafranski K, Xu Q, Tunggal B, Kummerfeld S, Madera M, Konfortov BA, Rivero F, Bankier AT, Lehmann R, Hamlin N, Davies R, Gaudet P, Fey P, Pilcher K, Chen G, Saunders D, Sodergren E, Davis P, Kerhornou A, Nie X, Hall N, Anjard C, Hemphill L, Bason N, Farbrother P, Desany B, Just E, Morio T, Rost R, Churcher C, Cooper J, Haydock S, van Driessche N, Cronin A, Goodhead I, Muzny D, Mourier T, Pain A, Lu M, Harper D, Lindsay R, Hauser H, James K, Quiles M, Madan Babu M, Saito T, Buchrieser C, Wardroper A, Felder M, Thangavelu M, Johnson D, Knights A, Loulseged H, Mungall K, Oliver K, Price C, Quail MA, Urushihara H, Hernandez J, Rabbinowitsch E, Steffen D, Sanders M, Ma J, Kohara Y, Sharp S, Simmonds M, Spiegler S, Tivey A, Sugano S, White B, Walker D, Woodward J, Winckler T, Tanaka Y, Shaulsky G, Schleicher M, Weinstock G, Rosenthal A, Cox EC, Chisholm RL, Gibbs R, Loomis WF, Platzer M, Kay RR, Williams J, Dear PH, Noegel AA, Barrell B, Kuspa A. 2005. The genome of the social amoeba Dictyostelium discoideum. Nature 435:43–57.

Elya C, Lok TC, Spencer QE, McCausland H, Martinez CC, Eisen M. 2018. Robust manipulation of the behavior of Drosophila melanogaster by a fungal pathogen in the laboratory. eLife 7:e34414.

Elya C, De Fine Licht HH. 2021. The genus Entomophthora: bringing the insect destroyers into the twenty-first century. IMA Fungus 12:34.

Fletcher K, Klosterman SJ, Derevnina L, Martin F, Bertier LD, Koike S, Reyes-Chin-Wo S, Mou B, Michelmore R. 2018. Comparative genomics of downy mildews reveals potential adaptations to biotrophy. BMC Genomics 19:851.

Floudas D, Binder M, Riley R, Barry K, Blanchette RA, Henrissat B, Martínez AT, Otillar R, Spatafora JW, Yadav JS, Aerts A, Benoit I, Boyd A, Carlson A, Copeland A, Coutinho PM, de Vries RP, Ferreira P, Findley K, Foster B, Gaskell J, Glotzer D, Górecki P, Heitman J, Hesse C, Hori C, Igarashi K, Jurgens JA, Kallen N, Kersten P, Kohler A, Kües U, Kumar TKA, Kuo A, LaButti K, Larrondo LF, Lindquist E, Ling A, Lombard V, Lucas S, Lundell T, Martin R, McLaughlin DJ, Morgenstern I, Morin E, Murat C, Nagy LG, Nolan M, Ohm RA, Patyshakuliyeva A, Rokas A, Ruiz-Dueñas FJ, Sabat G, Salamov A, Samejima M, Schmutz J, Slot JC, St. John F, Stenlid J, Sun H, Sun S, Syed K, Tsang A, Wiebenga A, Young D, Pisabarro A, Eastwood DC, Martin F, Cullen D, Grigoriev IV, Hibbett DS. 2012. The Paleozoic Origin of Enzymatic Lignin Decomposition Reconstructed from 31 Fungal Genomes. Science 336:1715–1719.

Galagan JE, Nusbaum C, Roy A, Endrizzi MG, Macdonald P, FitzHugh W, Calvo S, Engels R, Smirnov S, Atnoor D, Brown A, Allen N, Naylor J, Stange-Thomann N, DeArellano K, Johnson R, Linton L, McEwan P, McKernan K, Talamas J, Tirrell A, Ye W, Zimmer A, Barber RD, Cann I, Graham DE, Grahame DA, Guss AM, Hedderich R, Ingram-Smith C, Kuettner HC, Krzycki JA, Leigh JA, Li W, Liu J, Mukhopadhyay B, Reeve JN, Smith K, Springer TA, Umayam LA, White O, White RH, de Macario EC, Ferry JG, Jarrell KF, Jing H, Macario AJL, Paulsen I, Pritchett M, Sowers KR, Swanson RV, Zinder SH, Lander E, Metcalf WW, Birren B. 2002. The Genome of M. acetivorans Reveals Extensive Metabolic and Physiological Diversity. Genome Res 12:532–542.

Galagan JE, Calvo SE, Borkovich KA, Selker EU, Read ND, Jaffe D, FitzHugh W, Ma L-J, Smirnov S, Purcell S, Rehman B, Elkins T, Engels R, Wang S, Nielsen CB, Butler J, Endrizzi M, Qui D, Ianakiev P, Bell-Pedersen D, Nelson MA, Werner-Washburne M, Selitrennikoff CP, Kinsey JA, Braun EL, Zelter A, Schulte U, Kothe GO, Jedd G, Mewes W, Staben C, Marcotte E, Greenberg D, Roy A, Foley K, Naylor J, Stange-Thomann N, Barrett R, Gnerre S, Kamal M, Kamvysselis M, Mauceli E, Bielke C, Rudd S, Frishman D, Krystofova S, Rasmussen C, Metzenberg RL, Perkins DD, Kroken S, Cogoni C, Macino G, Catcheside D, Li W, Pratt RJ, Osmani SA, DeSouza CPC, Glass L, Orbach MJ, Berglund JA, Voelker R, Yarden O, Plamann M, Seiler S, Dunlap J, Radford A, Aramayo R, Natvig DO, Alex LA, Mannhaupt G, Ebbole DJ, Freitag M, Paulsen I, Sachs MS, Lander ES, Nusbaum C, Birren B. 2003. The genome sequence of the filamentous fungus Neurospora crassa. Nature 422:859–868.

Gandier J-A. 2017. Filling in the gaps of the spectrum-wide characterization of hydrophobins: An interface-active protein family with industrial potential. Toronto, Canada: University of Toronto.

Gandier J-A, Langelaan DN, Won A, O’Donnell K, Grondin JL, Spencer HL, Wong P, Tillier E, Yip C, Smith SP, Master ER. 2017. Characterization of a Basidiomycota hydrophobin reveals the structural basis for a high-similarity Class I subdivision. Sci Rep 7:45863.

Gandier J-A, Master E. 2018. Pichia pastoris is a Suitable Host for the Heterologous Expression of Predicted Class I and Class II Hydrophobins for Discovery, Study, and Application in Biotechnology. Microorganisms 6:3.

Gardner MJ, Hall N, Fung E, White O, Berriman M, Hyman RW, Carlton JM, Pain A, Nelson KE, Bowman S, Paulsen IT, James K, Eisen JA, Rutherford K, Salzberg SL, Craig A, Kyes S, Chan M-S, Nene V, Shallom SJ, Suh B, Peterson J, Angiuoli S, Pertea M, Allen J, Selengut J, Haft D, Mather MW, Vaidya AB, Martin DMA, Fairlamb AH, Fraunholz MJ, Roos DS, Ralph SA, McFadden GI, Cummings LM, Subramanian GM, Mungall C, Venter JC, Carucci DJ, Hoffman SL, Newbold C, Davis RW, Fraser CM, Barrell B. 2002. Genome sequence of the human malaria parasite Plasmodium falciparum. Nature 419:498–511.

Goffeau A, Barrell BG, Bussey H, Davis RW, Dujon B, Feldmann H, Galibert F, Hoheisel JD, Jacq C, Johnston M, Louis EJ, Mewes HW, Murakami Y, Philippsen P, Tettelin H, Oliver SG. 1996. Life with 6000 Genes. Science 274:546–567.

Haas BJ, Kamoun S, Zody MC, Jiang RHY, Handsaker RE, Cano LM, Grabherr M, Kodira CD, Raffaele S, Torto-Alalibo T, Bozkurt TO, Ah-Fong AMV, Alvarado L, Anderson VL, Armstrong MR, Avrova A, Baxter L, Beynon J, Boevink PC, Bollmann SR, Bos JIB, Bulone V, Cai G, Cakir C, Carrington JC, Chawner M, Conti L, Costanzo S, Ewan R, Fahlgren N, Fischbach MA, Fugelstad J, Gilroy EM, Gnerre S, Green PJ, Grenville-Briggs LJ, Griffith J, Grünwald NJ, Horn K, Horner NR, Hu C-H, Huitema E, Jeong D-H, Jones AME, Jones JDG, Jones RW, Karlsson EK, Kunjeti SG, Lamour K, Liu Z, Ma L, MacLean D, Chibucos MC, McDonald H, McWalters J, Meijer HJG, Morgan W, Morris PF, Munro CA, O’Neill K, Ospina-Giraldo M, Pinzón A, Pritchard L, Ramsahoye B, Ren Q, Restrepo S, Roy S, Sadanandom A, Savidor A, Schornack S, Schwartz DC, Schumann UD, Schwessinger B, Seyer L, Sharpe T, Silvar C, Song J, Studholme DJ, Sykes S, Thines M, van de Vondervoort PJI, Phuntumart V, Wawra S, Weide R, Win J, Young C, Zhou S, Fry W, Meyers BC, van West P, Ristaino J, Govers F, Birch PRJ, Whisson SC, Judelson HS, Nusbaum C. 2009. Genome sequence and analysis of the Irish potato famine pathogen Phytophthora infestans. Nature 461:393–398.

Hakanpää J, Paananen A, Askolin S, Nakari-Setälä T, Parkkinen T, Penttilä M, Linder MB, Rouvinen J. 2004. Atomic Resolution Structure of the HFBII Hydrophobin, a Self-assembling Amphiphile. Journal of Biological Chemistry 279:534–539.

Hakanpää J, Szilvay GR, Kaljunen H, Maksimainen M, Linder M, Rouvinen J. 2006. Two crystal structures of Trichoderma reesei hydrophobin HFBI--The structure of a protein amphiphile with and without detergent interaction. Protein Science 15:2129–2140.

Hanafy RA, Elshahed MS, Liggenstoffer AS, Griffith GW, Youssef NH. 2017. Pecoramyces ruminantium, gen. nov., sp. nov., an anaerobic gut fungus from the feces of cattle and sheep. Mycologia 109:231–243.

Hu X, Xiao G, Zheng P, Shang Y, Su Y, Zhang X, Liu X, Zhan S, St. Leger RJ, Wang C. 2014. Trajectory and genomic determinants of fungal-pathogen speciation and host adaptation. Proc Natl Acad Sci USA 111:16796–16801.

Jones L, Riaz S, Morales-Cruz A, Amrine KC, McGuire B, Gubler WD, Walker MA, Cantu D. 2014. Adaptive genomic structural variation in the grape powdery mildew pathogen, Erysiphe necator. BMC Genomics 15:1081.

Jones T, Federspiel NA, Chibana H, Dungan J, Kalman S, Magee BB, Newport G, Thorstenson YR, Agabian N, Magee PT, Davis RW, Scherer S. 2004. The diploid genome sequence of Candida albicans. Proc Natl Acad Sci USA 101:7329–7334.

Kämper J, Kahmann R, Bölker M, Ma L-J, Brefort T, Saville BJ, Banuett F, Kronstad JW, Gold SE, Müller O, Perlin MH, Wösten HAB, de Vries R, Ruiz-Herrera J, Reynaga-Peña CG, Snetselaar K, McCann M, Pérez-Martín J, Feldbrügge M, Basse CW, Steinberg G, Ibeas JI, Holloman W, Guzman P, Farman M, Stajich JE, Sentandreu R, González-Prieto JM, Kennell JC, Molina L, Schirawski J, Mendoza-Mendoza A, Greilinger D, Münch K, Rössel N, Scherer M, Vraneš M, Ladendorf O, Vincon V, Fuchs U, Sandrock B, Meng S, Ho ECH, Cahill MJ, Boyce KJ, Klose J, Klosterman SJ, Deelstra HJ, Ortiz-Castellanos L, Li W, Sanchez-Alonso P, Schreier PH, Häuser-Hahn I, Vaupel M, Koopmann E, Friedrich G, Voss H, Schlüter T, Margolis J, Platt D, Swimmer C, Gnirke A, Chen F, Vysotskaia V, Mannhaupt G, Güldener U, Münsterkötter M, Haase D, Oesterheld M, Mewes H-W, Mauceli EW, DeCaprio D, Wade CM, Butler J, Young S, Jaffe DB, Calvo S, Nusbaum C, Galagan J, Birren BW. 2006. Insights from the genome of the biotrophic fungal plant pathogen Ustilago maydis. Nature 444:97–101.

Kasson MT, Kasson LR, Wickert KL, Davis DD, Stajich JE. 2019. Genome Sequence of a Lethal Vascular Wilt Fungus, Verticillium nonalfalfae, a Biological Control Used Against the Invasive Ailanthus altissima. Microbiol Resour Announc 8:e01619–18.

Kasson MT, O’Donnell K, Rooney AP, Sink S, Ploetz RC, Ploetz JN, Konkol JL, Carrillo D, Freeman S, Mendel Z, Smith JA, Black AW, Hulcr J, Bateman C, Stefkova K, Campbell PR, Geering ADW, Dann EK, Eskalen A, Mohotti K, Short DPG, Aoki T, Fenstermacher KA, Davis DD, Geiser DM. 2013. An inordinate fondness for Fusarium: Phylogenetic diversity of fusaria cultivated by ambrosia beetles in the genus Euwallacea on avocado and other plant hosts. Fungal Genetics and Biology 56:147–157.

Katoh K, Standley DM. 2013. MAFFT Multiple Sequence Alignment Software Version 7: Improvements in Performance and Usability. Molecular Biology and Evolution 30:772–780.

Kazmierczak P, Kim DH, Turina M, Van Alfen NK. 2005. A Hydrophobin of the Chestnut Blight Fungus, Cryphonectria parasitica, Is Required for Stromal Pustule Eruption. Eukaryot Cell 4:931–936.

Kershaw MJ, Wakley G, Talbot NJ. 1998. Complementation of the Mpg1 mutant phenotype in Magnaporthe grisea reveals functional relationships between fungal hydrophobins. EMBO J 17:3838–3849.

Kim S, Ahn I-P, Rho H-S, Lee Y-H. 2005. MHP1, a Magnaporthe grisea hydrophobin gene, is required for fungal development and plant colonization: MHP1 of M. grisea for plant colonization. Molecular Microbiology 57:1224–1237.

Konkel Z, Scott K, Slot JC. 2021. Draft Genome Sequence of the Termite-Associated “ Cuckoo Fungus,” Athelia (Fibularhizoctonia) sp. TMB Strain TB5. Microbiol Resour Announc 10:e01230–20.

Kwan AHY, Winefield RD, Sunde M, Matthews JM, Haverkamp RG, Templeton MD, Mackay JP. 2006. Structural basis for rodlet assembly in fungal hydrophobins. Proc Natl Acad Sci USA 103:3621–3626.

Lauter FR, Russo VE, Yanofsky C. 1992. Developmental and light regulation of eas, the structural gene for the rodlet protein of Neurospora. Genes & Development 6:2373–2381.

Li W, Wang M, Bian X, Guo J, Cai L. 2016. A High-Level Fungal Diversity in the Intertidal Sediment of Chinese Seas Presents the Spatial Variation of Community Composition. Front Microbiol 7.

Lienemann M, Gandier J-A, Joensuu JJ, Iwanaga A, Takatsuji Y, Haruyama T, Master E, Tenkanen M, Linder MB. 2013. Structure-Function Relationships in Hydrophobins: Probing the Role of Charged Side Chains. Appl Environ Microbiol 79:5533–5538.

Linder MB. 2009. Hydrophobins: Proteins that self assemble at interfaces. Current Opinion in Colloid & Interface Science 14:356–363.

Littlejohn KA, Hooley P, Cox PW. 2012. Bioinformatics predicts diverse Aspergillus hydrophobins with novel properties. Food Hydrocolloids 27:503–516.

Loftus BJ, Fung E, Roncaglia P, Rowley D, Amedeo P, Bruno D, Vamathevan J, Miranda M, Anderson IJ, Fraser JA, Allen JE, Bosdet IE, Brent MR, Chiu R, Doering TL, Donlin MJ, D’Souza CA, Fox DS, Grinberg V, Fu J, Fukushima M, Haas BJ, Huang JC, Janbon G, Jones SJM, Koo HL, Krzywinski MI, Kwon-Chung JK, Lengeler KB, Maiti R, Marra MA, Marra RE, Mathewson CA, Mitchell TG, Pertea M, Riggs FR, Salzberg SL, Schein JE, Shvartsbeyn A, Shin H, Shumway M, Specht CA, Suh BB, Tenney A, Utterback TR, Wickes BL, Wortman JR, Wye NH, Kronstad JW, Lodge JK, Heitman J, Davis RW, Fraser CM, Hyman RW. 2005. The Genome of the Basidiomycetous Yeast and Human Pathogen Cryptococcus neoformans. Science 307:1321–1324.

Lovett, B. hydrophobin-survey, Zenodo (2022); doi.org/10.5281/zenodo.6981906

Lugones LG, Wösten HAB, Birkenkamp KU, Sjollema KA, Zagers J, Wessels JGH. 1999. Hydrophobins line air channels in fruiting bodies of Schizophyllum commune and Agaricus bisporus. Mycological Research 103:635–640.

Ma L, Chen Z, Huang DW, Kutty G, Ishihara M, Wang H, Abouelleil A, Bishop L, Davey E, Deng R, Deng X, Fan L, Fantoni G, Fitzgerald M, Gogineni E, Goldberg JM, Handley G, Hu X, Huber C, Jiao X, Jones K, Levin JZ, Liu Y, Macdonald P, Melnikov A, Raley C, Sassi M, Sherman BT, Song X, Sykes S, Tran B, Walsh L, Xia Y, Yang J, Young S, Zeng Q, Zheng X, Stephens R, Nusbaum C, Birren BW, Azadi P, Lempicki RA, Cuomo CA, Kovacs JA. 2016. Genome analysis of three Pneumocystis species reveals adaptation mechanisms to life exclusively in mammalian hosts. Nat Commun 7:10740.

Macias AM, Marek PE, Morrissey EM, Brewer MS, Short DPG, Stauder CM, Wickert KL, Berger MC, Metheny AM, Stajich JE, Boyce G, Rio RVM, Panaccione DG, Wong V, Jones TH, Kasson MT. 2019. Diversity and function of fungi associated with the fungivorous millipede, Brachycybe lecontii. Fungal Ecology 41:187–197.

Macindoe I, Kwan AH, Ren Q, Morris VK, Yang W, Mackay JP, Sunde M. 2012. Self-assembly of functional, amphipathic amyloid monolayers by the fungal hydrophobin EAS. Proc Natl Acad Sci USA 109.

McCabe PM, Van Alfen NK. 1999. Secretion of Cryparin, a Fungal Hydrophobin. Appl Environ Microbiol 65:5431–5435.

Mgbeahuruike AC, Kovalchuk A, Chen H, Ubhayasekera W, Asiegbu FO. 2013. Evolutionary analysis of hydrophobin gene family in two wood-degrading basidiomycetes, Phlebia brevispora and Heterobasidion annosums.1. BMC Evol Biol 13:240.

Miyauchi S, Kiss E, Kuo A, Drula E, Kohler A, Sánchez-García M, Morin E, Andreopoulos B, Barry KW, Bonito G, Buée M, Carver A, Chen C, Cichocki N, Clum A, Culley D, Crous PW, Fauchery L, Girlanda M, Hayes RD, Kéri Z, LaButti K, Lipzen A, Lombard V, Magnuson J, Maillard F, Murat C, Nolan M, Ohm RA, Pangilinan J, Pereira M de F, Perotto S, Peter M, Pfister S, Riley R, Sitrit Y, Stielow JB, Szöllősi G, Žifčáková L, Štursová M, Spatafora JW, Tedersoo L, Vaario L-M, Yamada A, Yan M, Wang P, Xu J, Bruns T, Baldrian P, Vilgalys R, Dunand C, Henrissat B, Grigoriev IV, Hibbett D, Nagy LG, Martin FM. 2020. Large-scale genome sequencing of mycorrhizal fungi provides insights into the early evolution of symbiotic traits. Nat Commun 11:5125.

Mondo Stephen J, Dannebaum RO, Kuo RC, Louie KB, Bewick AJ, LaButti K, Haridas S, Kuo A, Salamov A, Ahrendt SR, Lau R, Bowen BP, Lipzen A, Sullivan W, Andreopoulos BB, Clum A, Lindquist E, Daum C, Northen TR, Kunde-Ramamoorthy G, Schmitz RJ, Gryganskyi A, Culley D, Magnuson J, James TY, O’Malley MA, Stajich JE, Spatafora JW, Visel A, Grigoriev IV. 2017. Widespread adenine N6-methylation of active genes in fungi. Nat Genet 49:964–968.

Mondo Stephen J., Lastovetsky OA, Gaspar ML, Schwardt NH, Barber CC, Riley R, Sun H, Grigoriev IV, Pawlowska TE. 2017. Bacterial endosymbionts influence host sexuality and reveal reproductive genes of early divergent fungi. Nat Commun 8:1843.

Morris VK, Kwan AH, Mackay JP, Sunde M. 2012. Backbone and sidechain 1H, 13C and 15N chemical shift assignments of the hydrophobin DewA from Aspergillus nidulans. Biomol NMR Assign 6:83–86.

Morris VK, Kwan AH, Sunde M. 2013. Analysis of the Structure and Conformational States of DewA Gives Insight into the Assembly of the Fungal Hydrophobins. Journal of Molecular Biology 425:244–256.

Morris VK, Ren Q, Macindoe I, Kwan AH, Byrne N, Sunde M. 2011. Recruitment of Class I Hydrophobins to the Air:Water Interface Initiates a Multi-step Process of Functional Amyloid Formation. Journal of Biological Chemistry 286:15955–15963.

Mulder GH, Wessels JGH. 1986. Molecular cloning of RNAs differentially expressed in monokaryons and dikaryons ofSchizophyllum commune in relation to fruiting. Experimental Mycology 10:214–227.

Mycorrhizal Genomics Initiative Consortium, Kohler A, Kuo A, Nagy LG, Morin E, Barry KW, Buscot F, Canbäck B, Choi C, Cichocki N, Clum A, Colpaert J, Copeland A, Costa MD, Doré J, Floudas D, Gay G, Girlanda M, Henrissat B, Herrmann S, Hess J, Högberg N, Johansson T, Khouja H-R, LaButti K, Lahrmann U, Levasseur A, Lindquist EA, Lipzen A, Marmeisse R, Martino E, Murat C, Ngan CY, Nehls U, Plett JM, Pringle A, Ohm RA, Perotto S, Peter M, Riley R, Rineau F, Ruytinx J, Salamov A, Shah F, Sun H, Tarkka M, Tritt A, Veneault-Fourrey C, Zuccaro A, Tunlid A, Grigoriev IV, Hibbett DS, Martin F. 2015. Convergent losses of decay mechanisms and rapid turnover of symbiosis genes in mycorrhizal mutualists. Nat Genet 47:410–415.

Nagy LG, Ohm RA, Kovács GM, Floudas D, Riley R, Gácser A, Sipiczki M, Davis JM, Doty SL, de Hoog GS, Lang BF, Spatafora JW, Martin FM, Grigoriev IV, Hibbett DS. 2014. Latent homology and convergent regulatory evolution underlies the repeated emergence of yeasts. Nat Commun 5:4471.

Olsen KW. 1980. Internal residue criteria for predicting three-dimensional protein structures. Biochimica et Biophysica Acta (BBA) – Protein Structure 622:259–267. doi:10.1016/0005-2795(80)90036-7

Paananen A, Weich S, Szilvay GR, Leitner M, Tappura K, Ebner A. 2021. Quantifying biomolecular hydrophobicity: Single molecule force spectroscopy of class II hydrophobins. Journal of Biological Chemistry 296:100728.

Pendleton AL, Smith KE, Feau N, Martin FM, Grigoriev IV, Hamelin R, Nelson CD, Burleigh JG, Davis JM. 2014. Duplications and losses in gene families of rust pathogens highlight putative effectors. Front Plant Sci.

Peyretaillade E, Gonçalves O, Terrat S, Dugat-Bony E, Wincker P, Cornman RS, Evans JD, Delbac F, Peyret P. 2009. Identification of transcriptional signals in Encephalitozoon cuniculi widespread among Microsporidia phylum: support for accurate structural genome annotation. BMC Genomics 10:607.

Pham CLL, Rey A, Lo V, Soulès M, Ren Q, Meisl G, Knowles TPJ, Kwan AH, Sunde M. 2016. Self-assembly of MPG1, a hydrophobin protein from the rice blast fungus that forms functional amyloid coatings, occurs by a surface-driven mechanism. Sci Rep 6:25288.

Pham CLL, Rodríguez de Francisco B, Valsecchi I, Dazzoni R, Pillé A, Lo V, Ball SR, Cappai R, Wien F, Kwan AH, Guijarro JI, Sunde M. 2018. Probing Structural Changes during Self-assembly of Surface-Active Hydrophobin Proteins that Form Functional Amyloids in Fungi. Journal of Molecular Biology 430:3784–3801.

Pille A, Kwan AH, Cheung I, Hampsey M, Aimanianda V, Delepierre M, Latgé J-P, Sunde M, Guijarro JI. 2015. 1H, 13C and 15N resonance assignments of the RodA hydrophobin from the opportunistic pathogen Aspergillus fumigatus. Biomol NMR Assign 9:113–118.

Plett JM, Gibon J, Kohler A, Duffy K, Hoegger PJ, Velagapudi R, Han J, Kües U, Grigoriev IV, Martin F. 2012. Phylogenetic, genomic organization and expression analysis of hydrophobin genes in the ectomycorrhizal basidiomycete Laccaria bicolor. Fungal Genetics and Biology 49:199–209.

Quandt CA, Beaudet D, Corsaro D, Walochnik J, Michel R, Corradi N, James TY. 2017. The genome of an intranuclear parasite, Paramicrosporidium saccamoebae, reveals alternative adaptations to obligate intracellular parasitism. eLife 6:e29594.

R Core Team. 2021. A language and environment for statistical computing. R Foundation for Statistical Computing.

Raghukumar S. 2017. The Seagrass EcosystemFungi in Coastal and Oceanic Marine Ecosystems: Marine Fungi. Cham: Springer International Publishing. pp. 103–113.

Ren Q, Kwan AH, Sunde M. 2014. Solution structure and interface-driven self-assembly of NC2, a new member of the Class II hydrophobin proteins: Q. Ren et al. Proteins 82:990–1003.

Rineau F, Lmalem H, Ahren D, Shah F, Johansson T, Coninx L, Ruytinx J, Nguyen H, Grigoriev I, Kuo A, Kohler A, Morin E, Vangronsveld J, Martin F, Colpaert JV. 2017. Comparative genomics and expression levels of hydrophobins from eight mycorrhizal genomes. Mycorrhiza 27:383–396.

Rosenberg M, Kjelleberg S. 1986. Hydrophobic Interactions: Role in Bacterial Adhesion In: Marshall KC, editor. Advances in Microbial Ecology, Advances in Microbial Ecology. Boston, MA: Springer US. pp. 353–393.

RStudio Team. 2021. RStudio: Integrated Development Environment for R.

Russ C, Lang BF, Chen Z, Gujja S, Shea T, Zeng Q, Young S, Cuomo CA, Nusbaum C. 2016. Genome Sequence of Spizellomyces punctatus. Genome Announc 4:e00849–16.

Ryder LS, Cruz-Mireles N, Molinari C, Eisermann I, Eseola AB, Talbot NJ. 2022. The appressorium at a glance. Journal of Cell Science 135:jcs259857.

Scherrer S, De Vries OMH, Dudler R, Wessels JGH, Honegger R. 2000. Interfacial Self-Assembly of Fungal Hydrophobins of the Lichen-Forming Ascomycetes Xanthoria parietina and X. ectaneoides. Fungal Genetics and Biology 30:81–93.

Schmoll M, Tian C, Sun J, Tisch D, Glass NL. 2012. Unravelling the molecular basis for light modulated cellulase gene expression – the role of photoreceptors in Neurospora crassa. BMC Genomics 13:127.

Sharpton TJ, Stajich JE, Rounsley SD, Gardner MJ, Wortman JR, Jordar VS, Maiti R, Kodira CD, Neafsey DE, Zeng Q, Hung C-Y, McMahan C, Muszewska A, Grynberg M, Mandel MA, Kellner EM, Barker BM, Galgiani JN, Orbach MJ, Kirkland TN, Cole GT, Henn MR, Birren BW, Taylor JW. 2009. Comparative genomic analyses of the human fungal pathogens Coccidioides and their relatives. Genome Res 19:1722–1731.

Shin Y-K, Kim D-W, Lee S-W, Lee M-J, Gi Baek S, Lee T, Yun S-H. 2022. Functional roles of all five putative hydrophobin genes in growth, development, and secondary metabolism in Fusarium graminearum. Fungal Genetics and Biology 160:103683.

So K-K, Kim D-H. 2017. Role of MAPK Signaling Pathways in Regulating the Hydrophobin Cryparin in the Chestnut Blight Fungus Cryphonectria parasitica. Mycobiology 45:362–369.

St. Leger RJ, Staples RC, Roberts DW. 1992. Cloning and regulatory analysis of starvation-stress gene, ssgA, encoding a hydrophobin-like protein from the entomopathogenic fungus, Metarhizium anisopliae. Gene 120:119–124.

Stringer MA, Timberlake WE. 1995. dewA encodes a fungal hydrophobin component of the Aspergillus spore wall. Mol Microbiol 16:33–44.

Stringer MA, Timberlake WE. 1993. Cerato-Ulmin, a Toxin Involved in Dutch Elm Disease, Is a Fungal Hydrophobin. The Plant Cell 5:145.

Szilvay GR, Paananen A, Laurikainen K, Vuorimaa E, Lemmetyinen H, Peltonen J, Linder MB. 2007. Self-Assembled Hydrophobin Protein Films at the Air−Water Interface: Structural Analysis and Molecular Engineering. Biochemistry 46:2345–2354.

Tabima JF, Grünwald NJ. 2019. effectR: An Expandable R Package to Predict Candidate RxLR and CRN Effectors in Oomycetes Using Motif Searches. MPMI 32:1067–1076.

Terhem RB, van Kan JAL. 2014. Functional analysis of hydrophobin genes in sexual development of Botrytis cinerea. Fungal Genetics and Biology 71:42–51.

Tisserant E, Malbreil M, Kuo A, Kohler A, Symeonidi A, Balestrini R, Charron P, Duensing N, Frei dit Frey N, Gianinazzi-Pearson V, Gilbert LB, Handa Y, Herr JR, Hijri M, Koul R, Kawaguchi M, Krajinski F, Lammers PJ, Masclaux FG, Murat C, Morin E, Ndikumana S, Pagni M, Petitpierre D, Requena N, Rosikiewicz P, Riley R, Saito K, San Clemente H, Shapiro H, van Tuinen D, Bécard G, Bonfante P, Paszkowski U, Shachar-Hill YY, Tuskan GA, Young JPW, Sanders IR, Henrissat B, Rensing SA, Grigoriev IV, Corradi N, Roux C, Martin F. 2013. Genome of an arbuscular mycorrhizal fungus provides insight into the oldest plant symbiosis. Proc Natl Acad Sci USA 110:20117–20122.

Valsecchi I, Dupres V, Stephen-Victor E, Guijarro J, Gibbons J, Beau R, Bayry J, Coppee J-Y, Lafont F, Latgé J-P, Beauvais A. 2017. Role of Hydrophobins in Aspergillus fumigatus. JoF 4:2.

Valsecchi I, Stephen-Victor E, Wong SSW, Karnam A, Sunde M, Guijarro JI, Rodríguez de Francisco B, Krüger T, Kniemeyer O, Brown GD, Willment JA, Latgé J-P, Brakhage AA, Bayry J, Aimanianda V. 2020. The Role of RodA-Conserved Cysteine Residues in the Aspergillus fumigatus Conidial Surface Organization. JoF 6:151.

van Wetter M-A, Wosten HAB, Wessels JGH. 2000. SC3 and SC4 hydrophobins have distinct roles in formation of aerial structures in dikaryons of Schizophyllum commune. Mol Microbiol 36:201–210. doi:10.1046/j.1365-2958.2000.01848.x

Vandepol N, Liber J, Desirò A, Na H, Kennedy M, Barry K, Grigoriev IV, Miller AN, O’Donnell K, Stajich JE, Bonito G. 2020. Resolving the Mortierellaceae phylogeny through synthesis of multi-gene phylogenetics and phylogenomics. Fungal Diversity 104:267–289.

Viterbo A, Chet I. 2006. TasHyd1, a new hydrophobin gene from the biocontrol agent Trichoderma asperellum, is involved in plant root colonization. Molecular Plant Pathology 7:249–258.

Wessels JGH, de Vries OMH, Ásgeirsdóttir SA, Schuren FHJ. 1991a. Hydrophobin Genes lnvolved in Formation of Aerial Hyphae and Fruit Bodies in Schizophyllum. The Plant Cell 3:793–799.

Wessels JGH, De Vries OMH, Asgeirsdottir SA, Springer J. 1991b. The thn mutation of Schizophyllum commune, which suppresses formation of aerial hyphae, affects expression of the Sc3 hydrophobin gene. Journal of General Microbiology 2439–2445.

Wheeler TJ, Clements J, Finn RD. 2014. Skylign: a tool for creating informative, interactive logos representing sequence alignments and profile hidden Markov models. BMC Bioinformatics 15:7.

Will I, Das B, Trinh T, Brachmann A, Ohm RA, de Bekker C. 2020. Genetic Underpinnings of Host Manipulation by Ophiocordyceps as Revealed by Comparative Transcriptomics. Genes Genomes Genetics 10:2275–2296.

Winter DJ, Ganley ARD, Young CA, Liachko I, Schardl CL, Dupont P-Y, Berry D, Ram A, Scott B, Cox MP. 2018. Repeat elements organise 3D genome structure and mediate transcription in the filamentous fungus Epichloë festucae. PLoS Genet 14:e1007467.

Wösten HA, Schuren FH, Wessels JG. 1994. Interfacial self-assembly of a hydrophobin into an amphipathic protein membrane mediates fungal attachment to hydrophobic surfaces. The EMBO Journal 13:5848–5854.

Wösten HAB. 2001. Hydrophobins: Multipurpose Proteins. Annu Rev Microbiol 55:625–646.

Yoshimitsu Z, Nakajima A, Watanabe T, Hashimoto K. 2002. Effects of Surface Structure on the Hydrophobicity and Sliding Behavior of Water Droplets. Langmuir 18:5818–5822.

Youssef NH, Couger MB, Struchtemeyer CG, Liggenstoffer AS, Prade RA, Najar FZ, Atiyeh HK, Wilkins MR, Elshahed MS. 2013. The Genome of the Anaerobic Fungus Orpinomyces sp. Strain C1A Reveals the Unique Evolutionary History of a Remarkable Plant Biomass Degrader. Appl Environ Microbiol 79:4620–4634. doi:10.1128/AEM.00821-13

Yu L, Zhang B, Szilvay GR, Sun R, Jänis J, Wang Z, Feng S, Xu H, Linder MB, Qiao M. 2008. Protein HGFI from the edible mushroom Grifola frondosa is a novel 8 kDa class I hydrophobin that forms rodlets in compressed monolayers. Microbiology 154:1677–1685.

Zampieri F, Wösten HAB, Scholtmeijer K. 2010. Creating Surface Properties Using a Palette of Hydrophobins. Materials 3:4607–4625.

Zhang L, Villalon D, Sun Y, Kazmierczak P, van Alfen NK. 1994. Virus-associated down-regulation of the gene encoding cryparin, an abundant cell-surface protein from the chestnut blight fungus, Cryphonectria parasitica. Gene 139:59–64.

Zheng P, Xia Y, Xiao G, Xiong C, Hu X, Zhang S, Zheng H, Huang Y, Zhou Y, Wang S, Zhao G-P, Liu X, St Leger RJ, Wang C. 2011. Genome sequence of the insect pathogenic fungus Cordyceps militaris, a valued traditional chinese medicine. Genome Biol 12:R116.

